# Cell-type-specific coupling of single-unit spikes to cortical ripples conserved across macaque and human V1

**DOI:** 10.64898/2026.07.02.736041

**Authors:** Katarína Studeničová, Aitor Morales-Gregorio, Antonio Lozano, Xing Chen, Cristina Soto-Sánchez, Eduardo Fernandez, Paolo Papale, Karolína Korvasová

## Abstract

Cortical ripples are brief high-frequency (80–150 Hz) oscillatory bursts in the extracellular potential, associated with memory and attention. So far, it is unclear whether they are brain-state specific and which neuron types participate in their generation. We report cortical ripples in macaque V1 at rest, present with both eyes open and closed. Using massively parallel Utah array recordings, we identify six single-unit classes and describe their relation to ripple-band oscillations. The classes differ in firing-rate modulation, phase-locking strength, and preferred ripple-band phase. One class of putative deep-layer neurons exhibits non-sinusoidal phase locking, explained by a simple thresholding mechanism in a computational model. During ripples, most units fire irregularly with transient regular bursts. The ripple-associated spiking organisation is largely preserved during natural-image viewing and confirmed in V1 of a blind human. Together, these findings suggest cortical ripples arise from a shared circuit conserved across species and conditions.

**Graphical Abstract:** 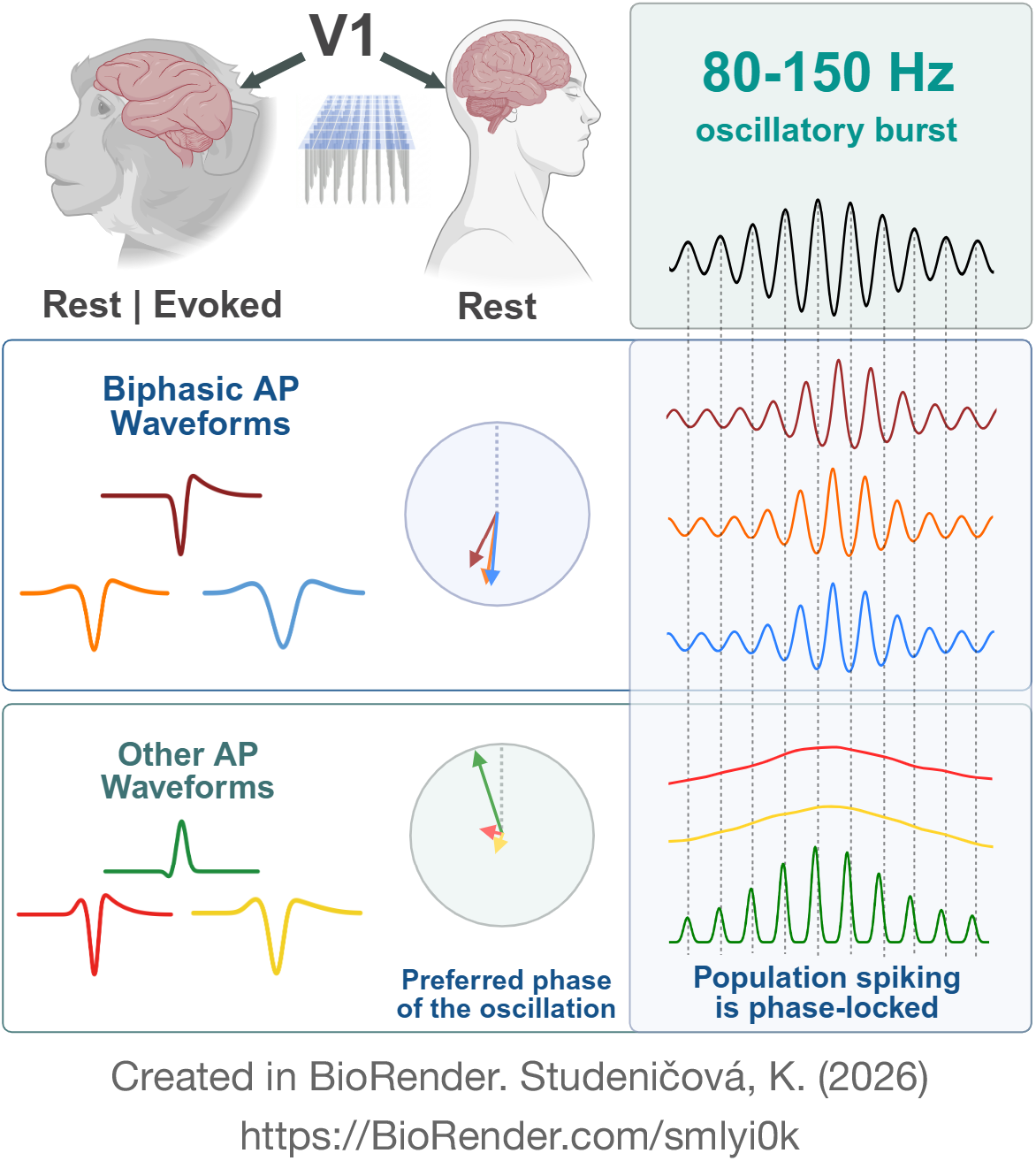

## Introduction

High-frequency oscillations (HFOs), neuronal rhythms spanning from 30 Hz all the way up to 500 Hz, reflect fast coordinated population activity in the cortex^1^. Unlike slow rhythms, HFOs typically emerge transiently and locally^1–4^. A subset of these events, characterized by brief bursts in the 80–150 Hz range and reminiscent of hippocampal ripples, has become known as “cortical ripples.”

Like their hippocampal counterparts, cortical ripples are thought to be involved in memory processes, showing an elevated rate in primary sensory areas during encoding and in higher-order areas during recall^5^. The ripple-like activity also increases with focused attention, correlating with faster detection times of behaviourally relevant events^6^. In rodents, cortical ripples have been identified in associative cortices during NREM sleep, with increased coupling to hippocampal ripples following learning^7^.

Complementing research on functional roles, resting-state recordings reveal spontaneous cortical ripple events. Early work in cats reported ripple-like activity in areas 5 and 7, in various states of vigilance and also in anesthesia^8^. In humans, short HFO bursts have been observed across widespread cortical regions, including primary sensory areas, during both sleep and wakefulness^5,9–12^. In non-human primates, however, cortical ripples have so far been described only in the context of visual attention^6^, and their occurrence and generating mechanisms during the resting state remain largely unexplored.

Although the number of studies about cortical ripples is growing, a detailed description of their generating mechanism is lacking. HFOs are generally associated with increases in local spiking activity^1,11,13^, pointing to the existence of a local circuit-like mechanism of HFO generation^9,14,15^. However, elucidating this mechanism requires identifying the specific cell types involved, characterising their precise spike timing relationships, and determining their laminar organisation within the cortical column.

Layer-dependent differences in high-frequency oscillations have been demonstrated across several studies^14–16^. Scheffer-Teixeira et al.^17^ showed that in the rat hippocampus, high-frequency activity in the pyramidal layer is strongly shaped by local spiking, whereas in superficial layers, transient increases in high-gamma power occur without accompanying spikes and reflect genuine oscillatory events. This pattern aligns with the findings of Leszczyński et al.^16^, who examined evoked HFOs in macaque primary visual cortex, primary auditory cortex, and human prefrontal cortex. According to their study, there is dissociation of HFOs and spikes in the superficial layers, but co-occurrence of oscillations with spiking activity in granular and deep layers.

Most human studies describe the involvement of putative excitatory and inhibitory units in HFOs but do not further resolve their cell-type identities^9,18^. In contrast, state-of-the-art spike-sorting applied in animal V1 reveals a far richer diversity of waveforms^19–21^. These studies show that both narrow- and broad-spiking units can be subdivided into multiple subclasses that occupy predominantly distinct cortical layers. Beyond the canonical negative-polarity waveforms of varying widths, attention has increasingly turned to triphasic units, characterized by a prominent pre-hyperpolarisation peak, and to units with predominantly positive-polarity waveforms. The biophysical origins of these waveform shapes remain debated, with proposed explanations including return currents and axonal signals^22^.

In this manuscript, we demonstrate the existence of spontaneously generated ripple-like bursts in the 80–150 Hz range in the primary visual cortex of macaque monkeys and blind humans. We identify six distinct classes of single units across putative granular and deep layers and characterize their contributions to HFO generation by analyzing phase-locking and firing-rate dynamics during cortical ripples. Together, these observations suggest the mechanism for cortical ripple generation could be an excitatory-inhibitory circuit present in Layer 4, with an additional rhythmic output in the deep layers. The responses during naturalistic visual stimulation suggest that the same oscillation-generating mechanism is active for both resting state and sensory processing. Moreover, the results are translatable to humans, as verified on two blind subjects with V1 Utah array implants.

## Results

### Brief bursts of elevated HFO activity appear in the resting state

Resting state activity was recorded from 3 macaque monkeys when seated in a room without visual stimulation and free to close their eyes. Single-unit activity and local field potentials (LFPs) were simultaneously acquired using 7–14 64-channel Utah arrays chronically implanted in V1 (Figure 1A). The frequency band of interest (80–150 Hz), referred to as the ripple band (RB), was selected based on existing literature, namely analyses of the macaque visual cortex recordings during visual attention^6^. High-frequency bursts lasting at least 40 ms within this band were detected using a dual-threshold algorithm^23^. For each channel, we bandpass filtered the signal within a frequency window of 80-150 Hz (sixth-order Butterworth filter) and calculated its Hilbert envelope. The envelope was further smoothed by a low-pass 40-Hz filter and normalized by its standard deviation. Next, we employed a double-threshold detection method with a minimal burst duration condition. Bursts were identified when the envelope exceeded an upper threshold of 3.5 standard deviations (SD), with duration defined as the interval during which the envelope remained above a lower threshold of 2.5 SD (Figure 1B). Events shorter than 40 ms were excluded. Detection was performed separately for eyes-open and eyes-closed periods; the latter likely encompassed multiple sleep stages (Figure 1C, Supplementary Figure 1A), which we did not differentiate further. We refer to the resulting events as spontaneous cortical ripples (S-CRs).

**Fig. 1.**
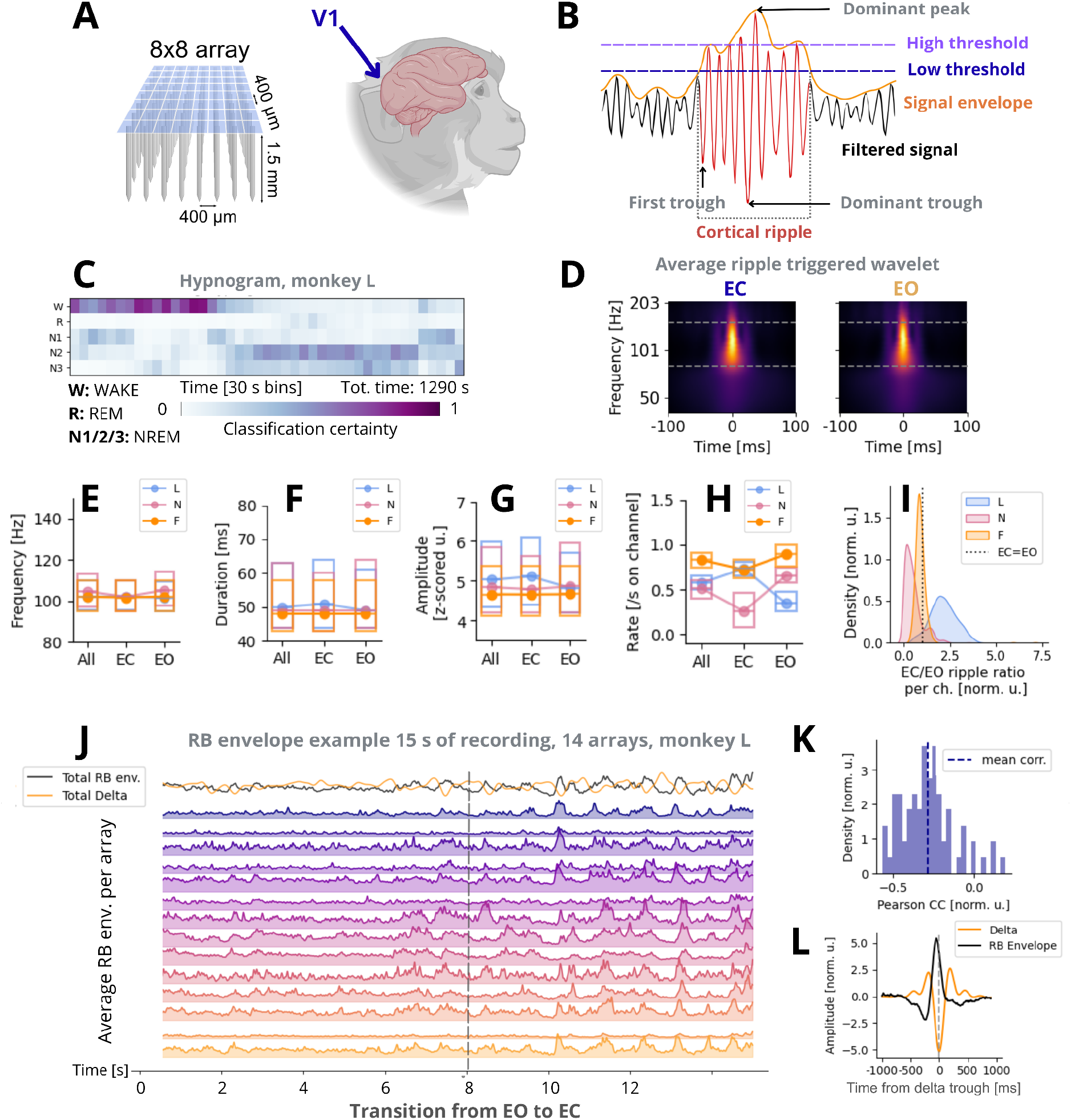
(S-CRs. Resting state.) **A.** Schematic of the Utah array used in the recordings (left), and the brain of the macaque monkey (right). “Recording Setup of Cortical Ripples Manuscript” created in BioRender. Studeničová, K. 2026 (https://BioRender.com/q0oxt4w). **B.** Schematics of the double threshold cortical ripple detection method. The 40-Hz lowpass Hilbert envelope of the filtered ripple band is thresholded in z-scored units. The time intervals above the lower threshold of at least 40 ms of duration with maximum amplitude above the higher threshold are considered as cortical ripples. **C.** Hypnogram of one of the recordings of monkey L. Each column corresponds to a classification result for a 30-s bin, based on YASA automated sleep scoring. All recording channels in V1 were used as input. For hypnograms of other recordings, see Supplementary Figure 1A. **D.** Ripple peak triggered wavelet spectrogram, eyes closed (EC) and open (EO). Rows are z-scored to compensate for the 1/f component. Plots for individual animals in Supplementary Figure 2A. **E, F, G.** The boxplots of the oscillation frequency of ripples, their duration, and amplitude, per animal, EC and EO, and pooled via the whole recording (All). One datapoint corresponds to one ripple. Boxplots display the median and the interquartile range. **H.** Number of ripples per second. One datapoint corresponds to one channel during one recording day. Eyes closed/open time intervals are considered separately. Boxplots display the median and the interquartile range. **I.** Distribution of the ratio of EC to EO ripple rate on individual channels, per animal. **J.** Average ripple band envelope on each of 14 V1 Utah arrays in monkey L, example 15 seconds of activity. Transition from EO to EC after the first 8 seconds. **K.** Histogram of Pearson correlation values between the average ripple band envelope on an array and the average of the delta (0-4 Hz) filtered signal on the same array. EC data only, all arrays, all animals. One data point represents a correlation value from the data on one array, one of the recording days. We tested whether the values are significantly negative with the Wilcoxon one-sided test: p<<10^-5^. **L.** The troughs of the average delta activity were used as triggers for the average delta-filtered signal on an array, and the average ripple band envelope on the same array. Plotted mean data 1 s around the trigger.

The detection procedure yielded oscillatory bursts that were well confined to the targeted high-frequency range (Figure 1D, E) and had a median duration of approximately 50 ms (Figure 1F). The median amplitude spanned values from 4.5 to 5.5 SD, varying slightly between subjects (Figure 1E-G). The number of bursts per second varied between the eyes-closed (EC) and eyes-open (EO) periods (Figure 1H, I). This effect was not consistent across monkeys, likely reflecting variability in brain state (Figure 1C; Supplementary Figure 1A), implantation site, and laminar localisation of electrode tips.

### Cortical ripples are associated with slow frequency phenomena

Ripple-like activity was accompanied by an LFP deflection followed by a prolonged upstate on the one-second timescale in both EC and EO conditions (Supplementary Figure 2B). At shorter timescales (∼100 ms), the averaged LFP signal revealed a regular ripple-like oscillation (Supplementary Figure 2C), indicating a high-frequency rhythm with stable cycle-to-cycle periodicity around the dominant peak^17^. During EC periods, the LFP exhibited an additional large-scale structure^8^, with synchronous global up- and down-states occurring in the delta frequency range, suggestive of transitions into sleep-related dynamics (Figure 1J). Motivated by this observation, we compared the 0–4 Hz delta band and the ripple band envelope, and found a clear anticorrelation (Figure 1J, top traces; Figure 1K, Wilcoxon one-sided test for Corr. values<0, p<<10^-5^). This relationship is further illustrated by the delta-trough-triggered ripple band envelope (Figure 1L), which demonstrates that ripple band activity peaks shortly before the delta trough and is attenuated during the 200–500 ms window surrounding it.

#### Six types of single-unit waveforms

We detected single neuron responses in all our recordings using custom data pre-processing and the state-of-the-art spike sorting algorithm KiloSort4^24^. Given the 400-µm spacing between electrodes in each Utah array, every unit was assigned to a single channel. In total, we identified 1608 single units across all the resting-state recording sessions, with some channels containing multiple units.

For each unit, we computed the average spike waveform, z-scored it, and extracted several waveform features (Figure 2A) to support subsequent classification. Specifically, we measured the trough-to-peak width (W), the height of the pre-hyperpolarisation peak (H), the amplitude from the mean to the maximum (UP), and the amplitude from the mean to the minimum (DOWN). First, we addressed the polarity of the waveform: Negative, if UP≤DOWN, or Positive if DOWN<UP (Pos). Negative-polarity waveforms were then subdivided into five classes. Based on width W, we defined Narrow (W ≤ 235 µs), Medium (235 µs < W ≤ 430 µs), and Wide (W> 430 µs) units (Figure 2B). Narrow units exhibited a bimodal distribution of the pre-hyperpolarisation peak height H (Figure 2C). Using a threshold of 1.2 SD, we further separated Narrow and Medium units into Biphasic (NarrBI, MedBI) when H was below the threshold, and Triphasic (NarrTRI, MedTRI) when H was above it. This procedure yielded six unit classes, which formed distinct clusters in t-SNE space (Figure 2D, E).

**Fig. 2.**
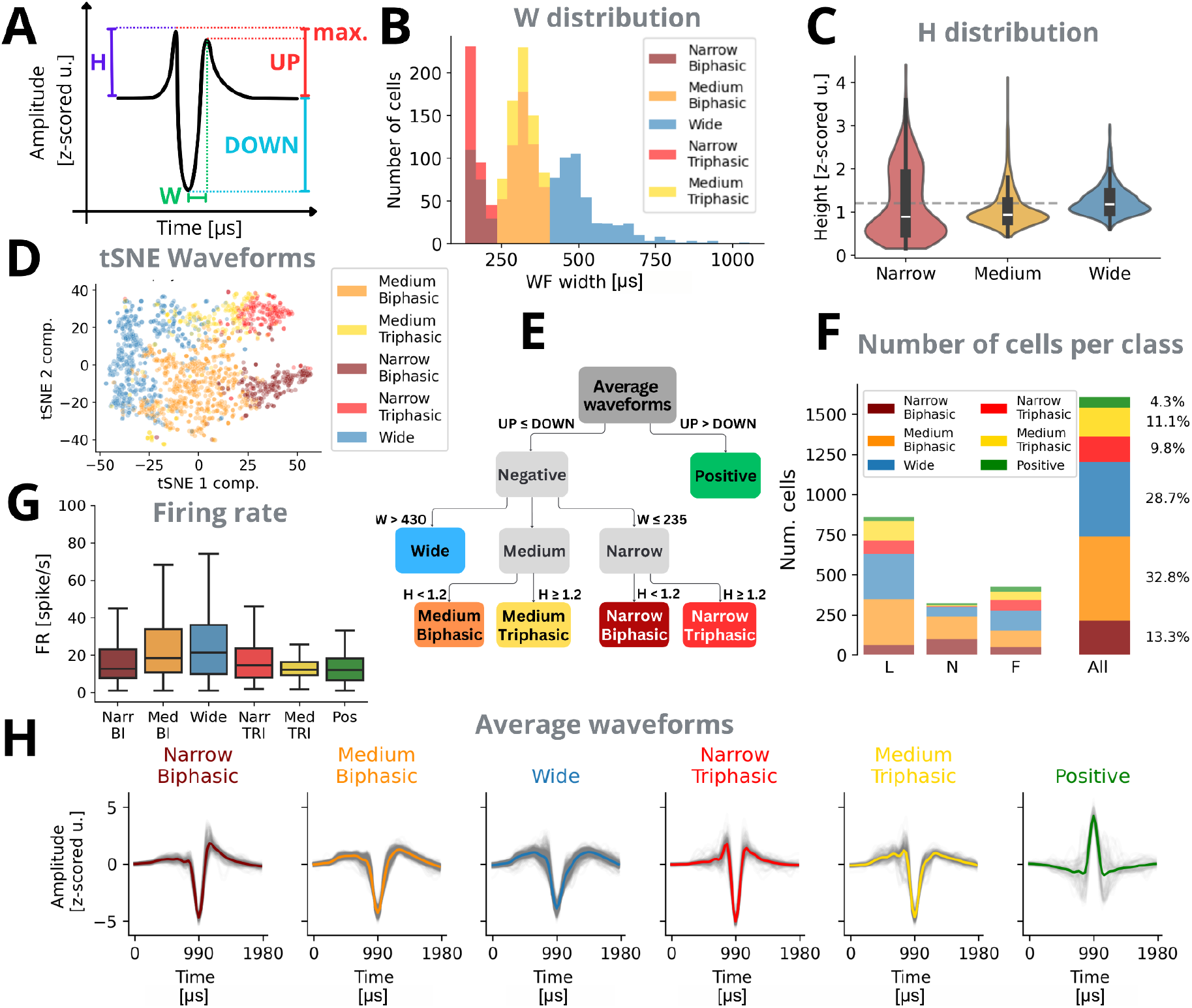
(SUA classification. Resting state.) **A.** Schematic of the measured properties of waveforms. The height of the first peak H, the waveform width W, and the amplitude in the negative direction DOWN. The amplitude in the positive direction UP is taken as the maximum of the height of both positive peaks of the waveform. **B.** Distribution of the waveform widths W, only negative dominant waveforms included. Color-coded based on the classification criteria, as depicted in E. **C.** Distribution of the first peak heights H per width class. **D.** An embedding of average single-unit waveforms on the first two t-SNE components. **E.** Unit classes definition flowchart. **F.** The number of single units per class detected, V1 pooled, per animal, and all animals pooled. **G.** Firing rate of single units. **H.** Average waveforms of single units per class are shown in gray, with the grand average for each unit type shown in color.

Most units exhibited Negative polarity (95.7%), with Medium Biphasic units being the most common (32.8%), followed by Wide (28.7%) and Narrow Biphasic (13.3%), see Figure 2F. Triphasic units accounted for ∼21% of the population. The distribution of unit types varied across monkeys, with monkey N almost completely lacking both Triphasic and Positive units, likely reflecting sampling differences and array placement. The firing rates of unit types differ systematically (Figure 2G). The average waveforms of units in each of the classes are shown in Figure 2H. Overall, the resulting classification aligns closely with recent single-unit characterisations in cat^19^, wallaby^20^, and macaque V1^21^, the latter recorded with Neuropixels.

### Phase locking differs systematically among unit types

When examining population activity across all Utah arrays in V1, we observed a clear co-occurrence between spiking events and elevations in the ripple band envelope (example in Figure 3A; for unit-class-resolved spiking, see Supplementary Figure 3A). The transition from eyes open (EO) to eyes closed (EC), when the brain state is changing towards an up/down state regime, was accompanied by changes in both single-unit spiking and transient fluctuations in ripple band power (Supplementary Figures 2C, 3A). During EC periods, the correlation between total spiking activity and the ripple band envelope increased compared with EO (Mann-Whitney test, p<0.001), indicating a tighter coupling between population firing and high-frequency activity in this state (Supplementary Figure 3B).

**Fig. 3.**
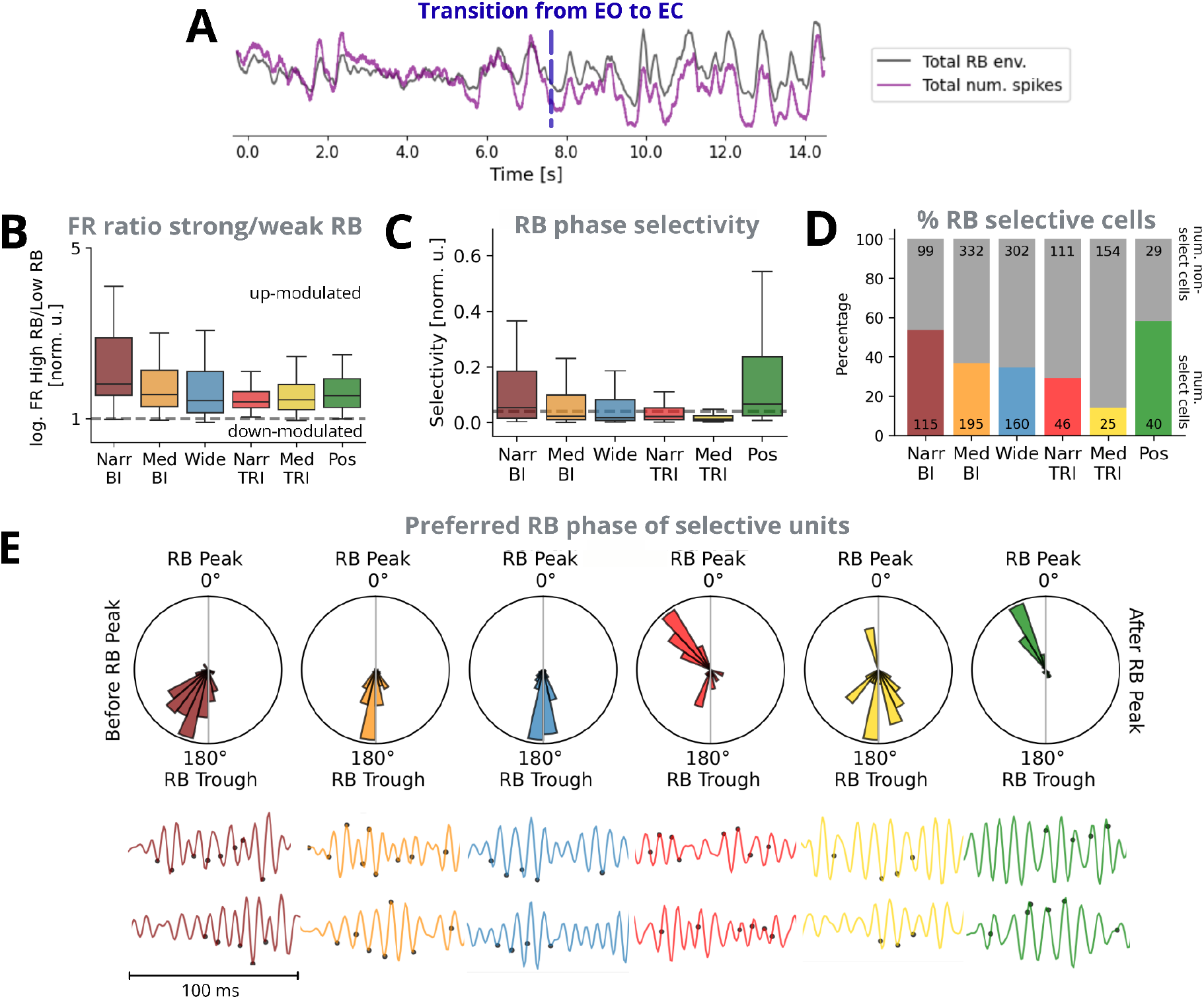
(Ripple band and spikes. Resting state.) **A.** Example 15s of activity, transition from EO to EC, monkey L, same time period as in Supplementary Figure 3A. The sum of the spiking activity shown as a purple line. The total RB envelope across all the channels in dark gray. Spiking activity per unit class in Supplementary Figure 3A. **B.** Ratio of firing rates during the RB envelope above its median value, and below its median value. Pooled per unit class, log-scale. A ratio>1 indicates up-modulation of spiking during high-amplitude ripple-band oscillation. **C.** Normalized RB phase selectivity for a unit class. Shuffle-based threshold 0.04 in dashed gray. **D.** Number and percentage of RB-phase-selective units, with selective units color-coded. Chance level <1%. **E.** Top: Histogram of preferred RB phases of selective units. One data point corresponds to a single unit of a given type. Bottom: Examples of 100ms of spiking activity of single-units superimposed on the RB filtered signal of a corresponding channel. Two units per class are shown. The filtered signal is color-coded, spikes are denoted as black dots. Plots B-E for individual animals are shown in Supplementary Figure 4. Comparison plots of panels B,C,E between EC and EO conditions are shown in Supplementary Figure 3C, D, E.

The observed degree of co-modulation between firing rate and the ripple band envelope moreover varied across unit types (Figure 3B), with a vast majority of units being up-modulated. The Biphasic units of a given width were up-modulated more strongly than their Triphasic counterparts (one-sided Mann-Whitney test, NarrBI>NarrTRI p-val<<0.001; MedBI>MedTRI p-val<<0.001). Overall, the highest positive modulation is present in Narrow Biphasic units (one-sided Mann-Whitney test with Bonferroni correction, NarrBI>Other for all 5 other classes, p<<0.001). Taken together, these responses reinforce the close, but unit-type-specific, relationship between HFOs and spiking activity^1,11,13^.

Next, we assessed the temporal precision of single-unit spiking relative to the phase of the ripple band signal. The instantaneous phase was obtained from the Hilbert transform of the 80–150 Hz band-pass-filtered LFP. For each unit, we computed a normalized selectivity (see STAR Methods), a number ranging from 0 to 1 quantifying the consistency with which spikes align to a particular oscillatory phase. A significance threshold (chance level <0.01) for this measure was determined using a shuffle-based procedure (100*Number of units shuffle data points, see STAR Methods). More than half of the Narrow Biphasic and more than 30% of Medium Biphasic and Wide units exhibited significant phase selectivity in RB (Figure 3C, D). Again, Triphasic units did not exhibit as high phase-selectivity as the same-width Biphasic classes (one-sided Mann-Whitney test, NarrBI>NarrTRI p-val<<0.001; MedBI>MedTRI p-val<<0.001). The comparatively weak modulation and phase-locking of Triphasic units suggests that not all neuronal classes contribute equally to HFO generation, and opens the question of the origin of Triphasic units, which we address in the Discussion. In contrast, units with Positive polarity waveforms displayed strong phase-locking, with more than half of the units meeting the significance criterion (Figure 3C, D).

Among the phase-selective units, the preferred oscillatory phase differed systematically across classes (Figure 3E). Wide units and Medium Biphasic units fired near the oscillatory trough, followed closely by Narrow Biphasic units. The narrow waveform shape and the phase shift of Narrow Biphasic units with respect to Wide units are indicative of a putative inhibitory nature of the Narrow Biphasic units, as we further comment on in Discussion^3,9^. Medium Triphasic units showed a similar, although less stable preference, also spiking near the trough. Narrow Triphasic units fired shortly before the oscillatory peak, preceding the Positive units.

In analyses of phase-locking and firing-rate modulation, we treated EC and EO oscillations as a single class of spontaneously occurring events. This approach emphasizes local circuit dynamics, while we acknowledge that large-scale network mechanisms shape ripple organisation differently across states. A direct comparison between EC and EO properties is provided in Supplementary Figure 3. Briefly, firing-rate modulation was stronger EC periods (Supplementary Figure 5C), whereas phase-locking characteristics remained consistent across EC and EO conditions (Supplementary Figures 3C, D).

### The ripple-generating process persists during natural-image viewing

Up to this point, we have characterized the relation between spiking organization and spontaneous cortical ripples during the resting state (S-CRs in RS). To extend these findings across visual conditions, we next examined activity during the natural-stimuli task.

The natural-image dataset (NATIM) was recorded by Papale et al.^25^ in monkeys N and F. Briefly, monkeys were shown images from the THINGS database^26^ (Figure 4A), each covering the receptive field. Each trial began with a fixation period. If fixation was successfully maintained, the trial continued after 300 ms with a sequence of four images, each presented for 200 ms and followed by a 200 ms gray screen. We analyzed the full recordings without reference to trial structure. We detected evoked cortical ripples (E-CRs) using the same methodology as was applied in the resting-state data. The amplitude, oscillation frequency, duration, and occurrence rate of E-CRs were comparable to those of spontaneous ripples in the resting state (Supplementary Figure 5, A-D).

**Fig. 4.**
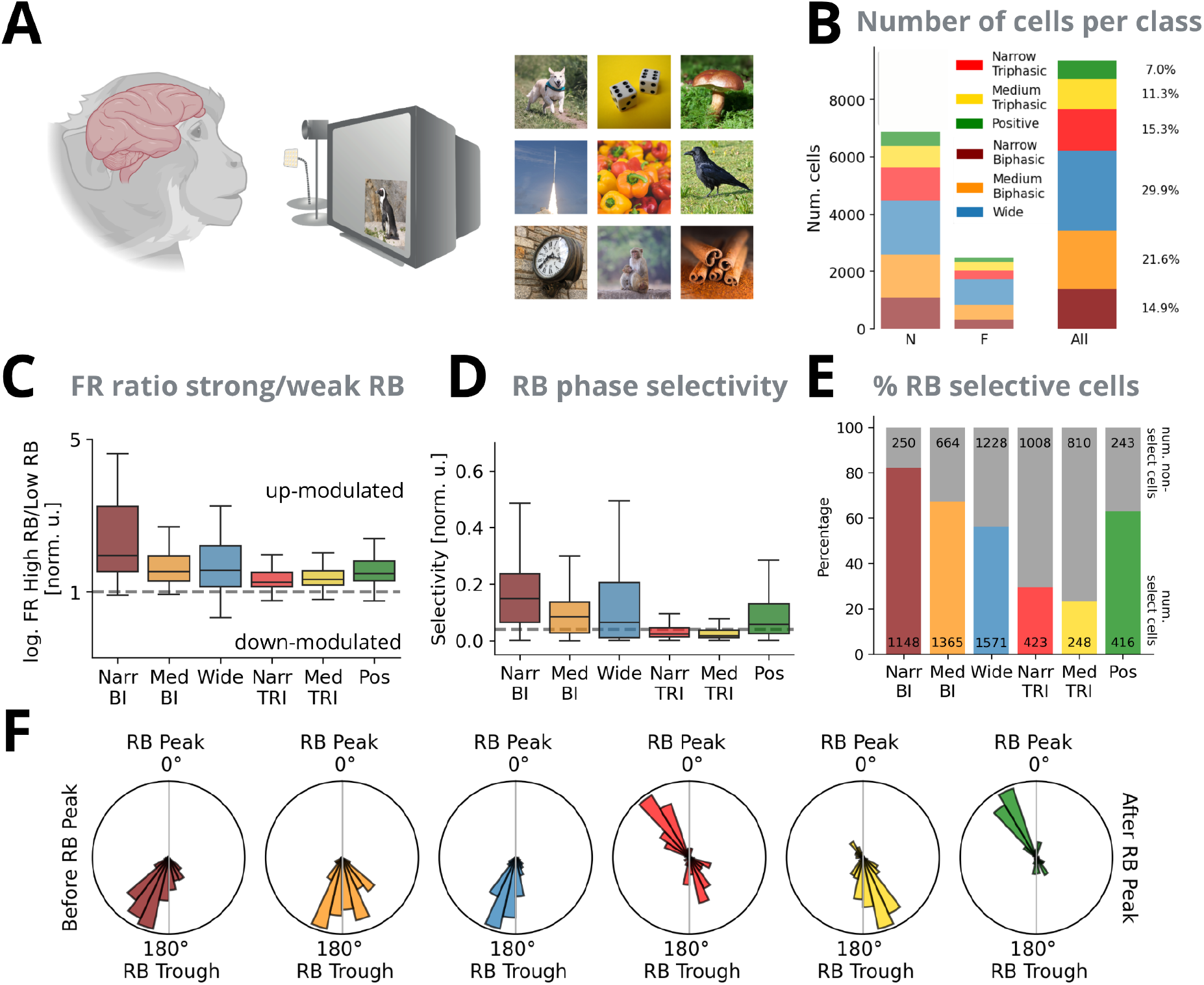
(Natural images.) The data are pooled across both animals, Natural images recordings only. References point to the analogous plot in the resting state analysis. **A.** Schematics of the Natural images stimulation paradigm. The monkey was shown images from the THINGS^26^ datasets, 200 ms per image. For details, see STAR Methods and Papale et al.^25^ “Recording Setup of Cortical Ripples Manuscript” created in BioRender. Studeničová, K. 2026 (https://BioRender.com/q0oxt4w) is licensed under CC BY 4.0. Freely publishable images from the THINGS dataset were used to create the illustration. **B.** The number of single units per class detected, V1 pooled, per animal, and all animals pooled. As in 2F. **C.** Ratio of firing rates during the RB envelope above its median value, and below its median value. Pooled per unit class, log-scale. As in 3B. **D.** Normalized RB phase selectivity for a unit class. Shuffle-based threshold in dashed gray. As in 3C. **E.** Number and percentage of RB phase-selective units. As in 3D. **F.** The histogram of preferred RB phases of selective units. One data point corresponds to one single unit of a given type. As in 3E.

Next, we examined how spiking activity contributed to the elevated ripple-band signal during natural images viewing. The waveform properties of the single units, such as the distributions of trough-to-peak widths and pre-hyperpolarisation peak heights, again revealed multiple distinct unit types. We classified these units using the same criteria applied in the resting state dataset (Figure 2E). Compared with RS (Figure 2F), the proportions of unit types in the population differed (Figure 4B): notably, Narrow Triphasic and Positive units were more prevalent in NATIM (RS VS NATIM % of unit type in population: POS 4.3% VS 7%, NarrTRI 9.8% VS 15.3%), particularly in monkey N. This likely reflects vertical shifts of the Utah arrays between RS and NATIM sessions. The total number of detected units was substantially higher in NATIM, when compared to RS, due to the larger number of recording sessions.

Among the six single-unit classes, the Biphasic units showed higher median up-modulation of spiking during periods of elevated ripple band power than the Triphasic units of the same width (Figure 4C), analogously to RS condition (Figure 3B). Again, analogously as in RS, the most pronounced up-modulation among all unit classes was visible in NarrBI units (Bonferroni-corrected Mann-Whitney one-sided test comparing NarrBI>Other. For NarrBI>WIDE p-val=7.8*10^-3^, for all other classes p-val<<10^-5^).

Next, we examined how neural activity during the eyes-open resting state (RS EO) compares with activity recorded in the NATIM protocol, in which the eyes remained open throughout the sessions. Apart from the triphasic units, all other classes (NarrBI, MedBI, Wide, Pos) were up-modulated more strongly in the NATIM condition (Supplementary Figure 5E, two-sided Mann-Whitney test, Bonferroni corrected p-vals: NarrBI, MedBI and Wide: p<<10^-5^, Pos: p=8.5*10^-3^).

Phase-selectivity values were significantly higher in NATIM than in RS EO for both Biphasic classes and for Wide units (Supplementary Figure 5F, two-sided Mann-Whitney test, Bonferroni corrected p-vals: p<<10^-5^ for all classes). Consequently, a larger proportion of units in the population of these classes were phase-selective in NATIM (Figure 4E) compared with RS (Figure 3D). Among Triphasic units, the phase selectivity was mostly below significance levels (Figure 4D, E), similarly as in RS (Figure 3C, D). For positive polarity units the phase selectivity of individual units was comparable in NATIM and in the RS EO (Supplementary Figure 5F). The preferred phases of the ripple band oscillation (Figure 4F) were consistent with the resting-state measurements (Figure 3E), indicating that although the modulation and phase-locking strength changed across conditions, the phase preference itself remained stable, possibly due to anatomical constraints on the circuitry.

### The unit classes are non-uniformly distributed across cortical space

The implanted Utah arrays spanned a substantial area of V1, providing broad horizontal coverage of the region (see figures depicting implantation positions in the original dataset publications^25,27^). The spatial distribution of unit types across this area was visibly non-uniform (e.g. see distribution for RS recordings in Supplementary Figure 6). To characterize this heterogeneity, we constructed a graph representing the probability that two units of a given type pair were detected on neighboring electrodes (Figure 5A, all RS and NATIM data pooled). A normalization was applied to highlight the unit-type pairs coupled beyond randomness (red edges in Figure 5A, see STAR Methods). This analysis revealed two distinct spatial groupings (highlighted in pale blue and pale orange in Figure 5A). The first grouping indicates that Positive and both Triphasic unit classes tend to occur in close spatial proximity. The second grouping comprises the Narrow and Medium Biphasic classes. The spatial distribution of the unit subgroups of one illustratory monkey is shown in Figure 5B (for other subjects see Supplementary Figure 7). Wide units did not show consistent spatial affinity with either grouping, being connected only to Pos, suggesting that this unit type is distributed broadly across the array, possibly comprising multiple subtypes with different spatial distributions.

**Fig. 5.**
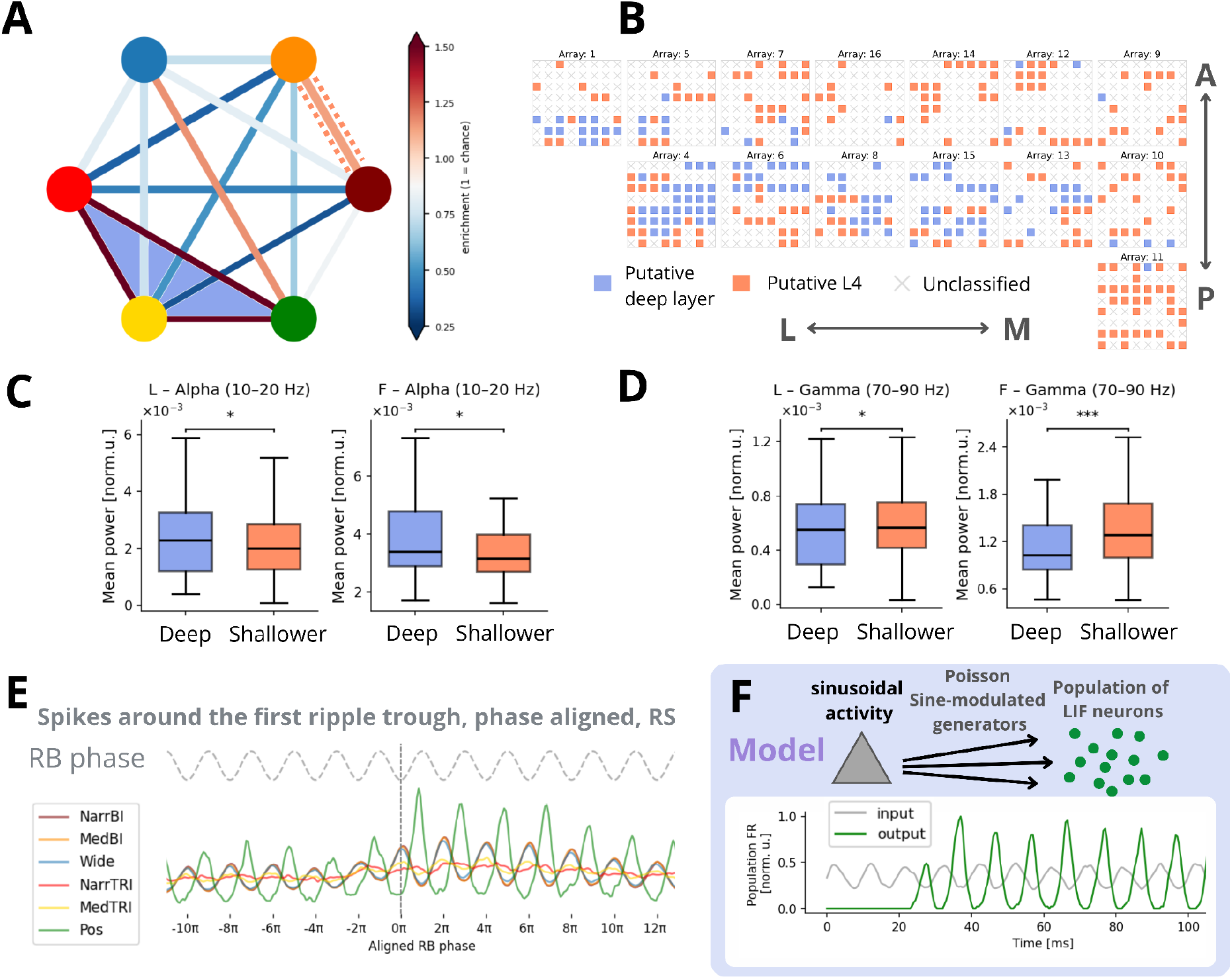
(Two subgroups of single units and model of oscillation generation.) **A.** Graph of neighbouring clustering of unit types. Each edge, color-coded, represents a normalised ratio of pairs of the given unit types detected on spatially close channels (normalization with respect to chance level, see STAR Methods). Unit types are colorcoded consistently with the rest of the manuscript. The two separated interconnected unit subgroups are highlighted in pale orange dotted lines and pale blue fill. **B.** For each channel, we assessed whether it contained more single units belonging to the blue-cluster unit group (NarrTRI+MedTRI+Pos) in panel A or the orange-cluster unit group (NarrBI+MedBI). Channels with no units, or with equal proportions of units from both cliques, were left unclassified. Data were pooled across all RS recordings for the monkey L. One gray box represents one Utah array, with 64 channels aligned in a square grid. The main orientation axes in the cortical space are depicted by arrows (Lateral-Medial, Anterior-Posterior, as in Figure 2 in the original dataset publication^27^). Distributions of unit groups for monkey N and F are shown in Supplementary Figure 7. **C.** Geometric means of power per channel in EO in the Alpha 10-20 Hz band, comparing putative deep-layer channels and putative shallower (granular layer) channels. Monkey L p-val. 1,5*10^-2^ and F p-val. 2,9*10^-2^, one-sided Mann–Whitney test. **D.** Same as C., but for the Gamma 70-90 Hz range. Monkey L p-val. 2,3*10^-3^ and F p-val. 6,9*10^-5^, one-sided Mann–Whitney test. **E.** The first ripple oscillatory trough triggered phase-aligned spikes. We aligned the spikes to the RB phase, and pooled the spiking activity around all the detected ripples, per unit class. Phase 0 corresponds to the trigger. **F.** Schema of the model (top). The first 100 ms of the model activity (bottom). Sine-modulated Poisson spiking in gray. Resulting activation of the LIF population in green.

**Fig. 6.**
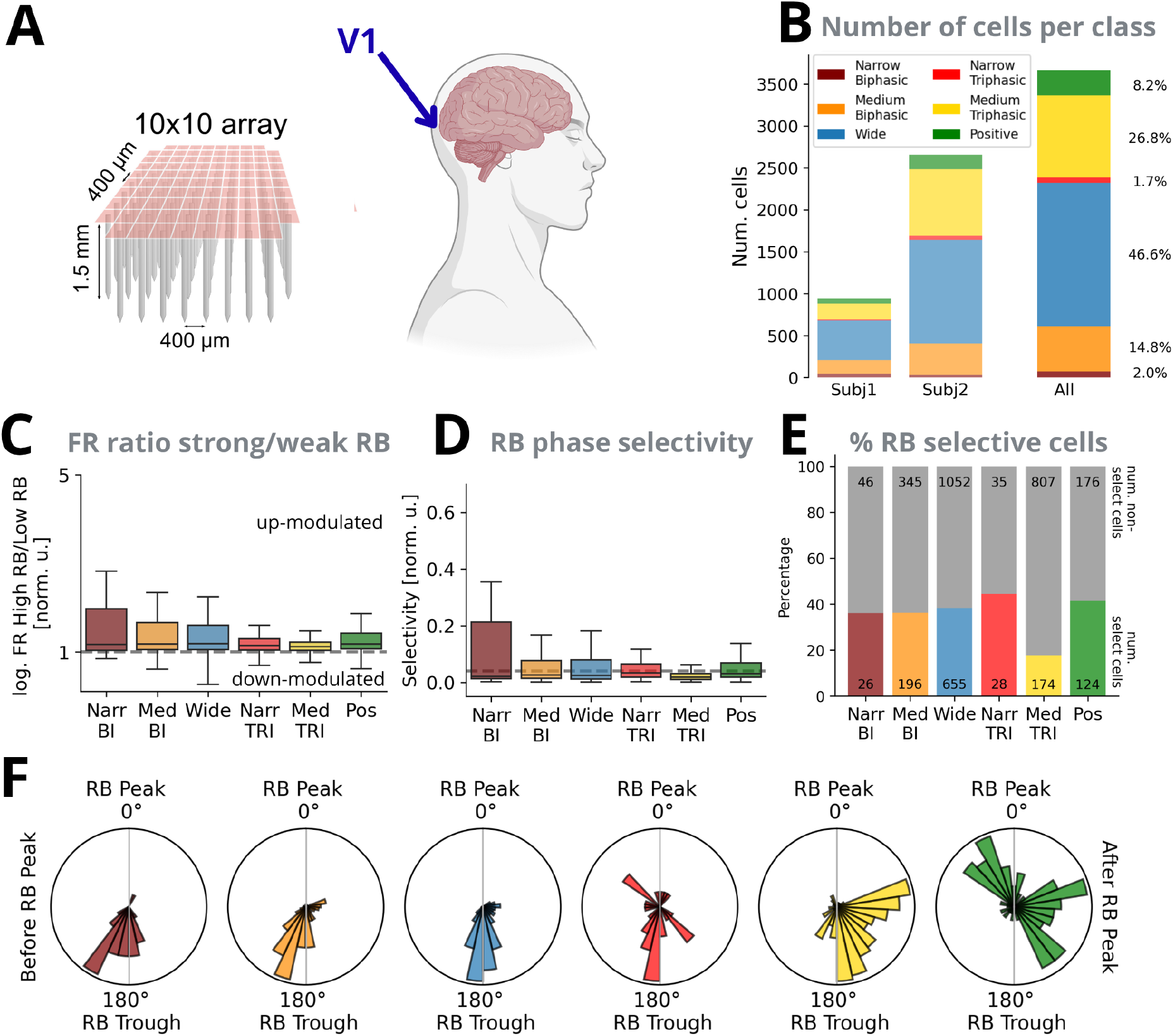
(Blind human.) References point to the analogous plot in the macaque resting state analysis. **A.** Schematic of the Utah array implant used in the human recordings (left), and the brain of the human subject (right). “Recording Setup of Cortical Ripples Manuscript” created in BioRender. Studeničová, K. 2026 (https://BioRender.com/q0oxt4w) is licensed under CC BY 4.0. **B.** The number of single units per class detected. Individual bar plots are shown per subject, and for the pooled data. As in 2F. **C.** Ratio of firing rates during the RB envelope above its median value, and below its median value. Pooled per unit class across both subjects, log-scale. As in 3B. **D.** Normalized RB phase selectivity for a unit class, pooled across both subjects. Shuffle-based threshold in dashed gray. As in 3C. **E.** Number and a ratio of RB phase selective units. As in 3D. **F.** The histogram of preferred RB phases of selective units. One data point corresponds to one single unit of a given type. As in 3E. Comparative statistics of panels C-F per subject are shown in Supplementary Figure 10.

Given the columnar and laminar organization of V1^28^, we hypothesize that the observed co-occurrence patterns arise primarily from small vertical differences in electrode depth, rather than from lateral inhomogeneities within a single layer. Consistent with this interpretation, Neuropixels recordings^21^ demonstrate that Positive-polarity units are most abundant in the deep layers and white matter. Among negative-polarity units, Triphasic and Wide waveforms are the most common in the deep layers. Thus, we consider the clustered Triphasic and Positive classes as putative deep-layer units. The remaining clustered unit types — Narrow Biphasic, Medium Biphasic (orange subgroup in Figure 5A, B) — likely reflect populations predominantly located shallower, we suppose Layer 4.

This putative laminar assignment can be further validated by spectral analyses (Figure 5C, D; Supplementary Figure 8A,B) based on Mendoza-Halliday et al.^14^ According to this study, we hypothesize that the spectral power in Alpha band (10-20 Hz) will be more pronounced on the channels where the deep-layer units are located, compared to putative granular-layer units. On the contrary, the Gamma band (70-90 Hz) will show opposite gradient, being stronger on the shallower channels. We confirmed this both in RS EO (Figure 5C, D), and in fixation periods in the NATIM condition (Supplementary Figure 8A, B). In the spectral comparison for RS only the monkeys L and F were included, since the monkey N did not have enough blue-group channels. For a full argument on putative layer assignment see Discussion.

### Unit subgroups synchronise to support oscillatory dynamics

The spiking activity locked to the oscillation may exhibit two forms of synchrony^29^: either individual units fire regularly and in phase, or units fire irregularly, but the population as a whole remains phase-synchronized. Overall, the unit activity was highly variable across the population and time, suggesting the synchronous-irregular firing regime. Nevertheless, we observed transient episodes in which single units fired repeatedly at a consistent phase of the ripple band oscillation (Figure 3E bottom), indicative of the regular-firing regime occurring at least transiently. Examples from two units of each class illustrate this behaviour, with each unit reliably spiking at its preferred phase, consistent with Figure 3E, top plots. Therefore, we were not able to reliably distinguish between regular and irregular regimes, and we further investigated population-wide locking effects without assuming either of these.

To describe the time course of unit firing during each ripple, we analyzed single-unit spiking activity time-locked to individual S-CRs (Figure 5E). For each S-CR, the single-unit activity recorded on the corresponding channel was aligned to the oscillatory phase, with phase zero defined at the first trough of the ripple burst (as illustrated in Figure 1B). During each burst, which was typically comprised of at least four oscillatory cycles, Biphasic, Wide and Positive waveform populations exhibited elevated rhythmic spiking (Figure 5E), consistent with the rate modulation previously assessed in Figure 3B. At the ripple onset, the Biphasic and Wide units activated first, followed by the Positive units. The phase-locking profiles of the Biphasic and Wide units displayed a clear sinusoidal structure, aligning with expectations for activity generated by a local excitatory–inhibitory circuit^3,9^. On the other hand, the Positive units exhibited sharp, non-sinusoidal activation, in line with the previously observed strong phase-locking (Figure 3C, D).

To provide a plausible explanation of the observed phase-difference and strong non-sinusoidal locking of Positive units, we constructed a simple computational model in the NEST simulator^30^ (Figure 5F). The model comprised a population of 2,500 alpha-function Leaky Integrate-and-Fire (LIF) neurons without mutual connections. Inputs to this population approximated the sine-like rhythmic drive originating from the assumed granular-layer units and were implemented using Poisson spike-train generators with sinusoidal modulation. Multiple Poisson generators operating at 100 Hz, each with a slight phase offset, were randomly connected to the model neurons with standard connection parameters (STAR Methods). The model operated in a synchronous-irregular regime^29^ with a mean firing rate of ∼9 Hz, in line with the data (Figure 5E). The resulting spiking activity (Figure 5F) reproduced the dynamics observed in Positive units during cortical ripples (Figure 5E), particularly across the four oscillatory cycles following the trigger. Model neurons fired with a phase delay of nearly half an oscillatory cycle relative to the granular-layer input, consistent with the phase relationships measured in the data (Figure 5E). They also displayed low baseline firing with sharp rhythmic increases, mirroring the real data patterns in Figure 5E. These results suggest that the non-sinusoidal rhythmic activation and strong phase-locking of Positive units can arise from a combination of sinusoidal shallower layer input together with a thresholding mechanism of the postsynaptic neurons.

In NATIM condition, when examining the phase-aligned population spiking triggered by E-CRs (Supplementary Figure 8C), the Biphasic and Wide units again exhibited a clear sinusoidal modulation, closely resembling the pattern observed during spontaneous ripples (Figure 5E). Positive-polarity units showed noisier dynamics, not showing as pronounced sharp activity peaks (Supplementary Figure 8C). Triphasic units contributed only weakly, not showing a rhythmic component, in a similar manner as in the resting state.

### Oscillatory-coupled spike activation is reproduced in blind human V1

We analyzed resting-state recordings from two volunteers with acquired blindness implanted with a 96-channel Utah array in V1 (STAR Methods). The volunteers sat in the experimental room without additional tasks or stimuli. The dataset comprises morning and evening sessions; in the intervening periods, additional recordings and electrical stimulation experiments were conducted but are not analyzed here. Data preprocessing, spike sorting, ripple detection, and unit classification were performed following the same pipeline as in the macaque monkeys (STAR Methods), allowing us to directly assess the translatability of the approach to the human cortex.

We first detected cortical ripples. The resulting ripples had a rate, duration, frequency, and amplitude (Supplementary Figure 9A-D) similar to those observed in macaque monkeys (Figure 1E-H). We then classified unit waveforms using the same criteria as in macaques (Figure 2E). The resulting proportions of unit types differed from those in monkeys, most notably with fewer narrow units and more wide units; this discrepancy is likely due to a suboptimal width-classification threshold (Supplementary Figure 9E). Nonetheless, all six unit types identified in monkeys were also present in the human single-unit population.

As in monkeys, most of the units had firing rates elevated during the periods of strong RB activity (Figure 6C). However, the Narrow Biphasic did not show median modulation higher than other classes, as in the case of monkeys (Figure 3B, 4C). Importantly, the preferred ripple-band phase of each unit type (Figure 6F) corresponded with the monkey results (Figure 3E, 4F), although being less stable. The biphasic and wide units locked to the phase near the trough, with narrow units shortly following the medium and wide units. A subset of positive units locked to the phase near the peak as in the monkey datasets, although overall the preference of this unit class was much noisier. The triphasic units also showed noisier phase locking to multiple oscillatory phases.

In the phase-aligned, ripple-triggered spiking diagram (Supplementary Figure 9F), the overall dynamics was noisier, but not contradicting analogous analyses in monkeys (Figure 5E, Supplementary Figure 8C). The results were mostly consistent across both subjects (Supplementary Figure 10). Overall, although the width-based classification threshold may require human-specific tuning, the coupling between unit type and ripple-band activity is reproducible in the human cortex.

## Discussion

Our findings demonstrate spontaneously occurring ripple-like bursts in macaque and human V1, in the 80–150-Hz range. We distinguished six classes of single units, outlining how they may contribute to these oscillatory events in the early visual cortex, both during spontaneous activity and viewing of natural images. Our results suggest the mechanism for ripple generation might be an excitatory–inhibitory circuit, likely localised in Layer 4. We also observed a non-sinusoidal deep-layer rhythmic pattern, possibly arising from additional thresholding of shallower layers outputs, as demonstrated by a simple computational model.

As for the putative Layer 4 populations expressing the sinusoidal rhythmicity, we were not able to further verify the inhibitory or excitatory nature of the measured units. However, the observed difference in the preferred RB phase (Figure 3E) between Narrow Biphasic and Wide units, corresponds well to the putative inhibitory and excitatory nature of NarrBI and Wide units, respectively, reflecting inhibition stabilising the oscillation^3,9^. The phase difference between NarrBI and Wide units was also present in humans (Figure 6F). Both observed differences are in the range of 1-2ms, consistent with monosynaptically interconnected excitatory-inhibitory circuit. The interspecies differences can arise from the differences in cortical architecture of non-human primates and humans^31^. Otherwise, these results are remarkably consistent both intra- and interspecies.

Within the putative Deep-layer group identified in Figure 5A (Positive+ Triphasic), only the Positive units exhibited strong phase-locking, which was moreover phase shifted with respect to the putative Layer 4 units. This observation supports the hypothesis that rhythmic firing in the deep layers is not generated by a local excitatory–inhibitory circuit but is instead broadcast from other layers. We did not observe LFP polarity reversal across recording channels, thus we estimate that the phase shift represents the time difference of ∼5ms between the sine-generating population firing and the Positive units. This observed phase lag is consistent with monosynaptic transmission.

Altogether, we demonstrated the plausibility of the scenario where the putative Layer 4 excitatory-inhibitory circuit broadcasts the ripple-band oscillation to deeper layers. Elucidating the generation mechanisms of cortical ripples is crucial for a full explanation of their involvement in information integration and memory processes in the neocortex.

### Spike leaking or a true oscillation?

A substantial body of literature reports that high-frequency activity co-occurs with elevated spiking^11,13,16,17^, raising the question of whether such spike-associated HFOs reflect genuine oscillatory events or merely broadband consequences of population firing. Although we acknowledge the ongoing challenges in defining HFOs rigorously^16,17^, our data provide several indications of true oscillatory structure. First, the ripple-triggered spectrogram (Figure 1D) reveals power that is well confined to the targeted frequency band, consistent with a narrowband rhythm rather than broadband leakage. Second, the precise millisecond-scale spike timing observed in putative Layer 4 units supports the presence of an intrinsic oscillation-generating circuit capable of organizing firing with high temporal precision. In the deeper layers, where positive-polarity units likely broadcast an inherited rhythm, the phenomenon may indeed approach the non-oscillatory spike-leak interpretation^13,17^. Together, these observations suggest that while some high-frequency events may reflect spike-related broadband activity, a substantial subset exhibits genuine oscillatory dynamics.

### Layer identification

Addressing the question of laminar distribution of detected units, a recent study using Neuropixels provides a unique view of a laminar organisation of waveforms in the cortex of macaque monkeys^21^. Importantly, a high number of units with positive waveforms was observed in the deep layers and white matter. Other types of units detected in the deep layers were predominantly those that correspond to our Wide and Triphasic classes. Based on this, we concluded that the spatial clustering of Triphasic + Positive units in our analysis (Figure 5A, blue subgroup) was likely due to the electrode tip reaching the deep cortical layers.

Another cluster of unit types we identified consisted of Biphasic Narrow and Biphasic Medium (Figure 5A, orange subgroup). Since this subgroup of units was rarely co-detected with Triphasic and Positive waveforms, we posit that it was not localized in the deep layers. We further posit that the recording was not from the shallow layers either, for three reasons. Firstly, we generally expect only a small number of Narrow Biphasic units to be detectable in the shallow layers^21^. Secondly, given the length of the Utah array shanks (1.5 mm) and thickness of layers in the macaque V1^31^, the electrodes were long enough to reach at least Layer 4 ^25,27^, although note that the recording locations were not conclusively verified histologically. Lastly, multiple Utah arrays were split into two parts of neighbouring channel types, recording one of the unit groups on one part of the electrodes, and the other group on the other part of the channels (Supplementary Figure 7). This implies that both co-detected groups are located in the neighbouring laminae. Finally, the aforementioned study^21^ agrees that the Narrow Biphasic waveforms are mostly localized in Layer 4. Therefore, we conclude that the group of co-detected unit types of Biphasic Narrow and Medium is localized in Layer 4.

To further examine our hypothesis that the distinct groups of units were recorded in different laminae, we analysed the modulations of alpha and gamma power. A recent study by Mendoza-Halliday et al.^14^ proposed that putative deep-layer channels should exhibit dominant alpha-band activity, whereas shallower channels show more pronounced gamma power. We observed this pattern consistently, both during eyes-open rest and during the gaze fixation period (Figure 5C, D, Supplementary Figure 8A, B).

The spatial clustering of unit types did not seem to be driven by the horizontal position of the Utah arrays within the V1 area (Supplementary Figures 6, 7), although the clustering of the blue channels across neighbouring arrays might reflect cortical curvature. Overall, the Utah arrays may have either been implanted in a slightly slanted angle with respect to the cortical layers, or in the place of curved laminae, thus yielding recordings from distinct layers, presumably 4 and 5/6. Furthermore, since the recordings were chronic, the arrays might have shifted slightly vertically, increasing the variability across days. Additional effects might have been caused by gradual lesioning of the cortex. A limitation of our approach is that we were unable to further validate our conclusions with histological verification of electrode tip locations, due to chronic lesioning of the tissue^25,27^.

### Origin of non-standard units

Although waveform width is typically emphasized as the primary criterion for single-unit classification, our analyses revealed additional distinctions based on the amplitude of the pre-hyperpolarization peak. Triphasic waveforms of this kind have been reported previously in high-density microelectrode recordings from cell cultures^32–35^, as well as in vivo, both in humans^36,37^, and in other species^19–22,38^. Notably, muscimol-induced cortical silencing does not attenuate their spiking^19,38^, in contrast to the spiking of biphasic units. This observation has led to the hypothesis that these waveforms arise from a non-somatic origin, most plausibly axonal. Proposed mechanisms include axonal forward propagation, return currents, or dendritic backpropagation^19,22,38^. Although the Utah array does not permit definitive identification of the recorded cellular compartment, the weak entrainment of these units during high-frequency oscillations, together with their co-occurrence with Positive units in deep layers, supports the interpretation that they reflect a likely axonal source consistent with prior reports.

Positive units, which are observed predominantly in the deep layers and white matter^21^, are widely interpreted as reflecting axonal signals^19–22,39^. Although our data show that these units participate in ripple-band activity, a simple computational model suggests that they may not constitute part of the excitatory–inhibitory circuit generating the rhythm. Instead, they may inherit the broadcast oscillation while being recorded at an axonal locus. The available evidence does not allow us to determine whether these axons originate from thalamic or cortical sources.

Overall, while spike-waveform width has historically served as the dominant criterion for single-unit classification, our findings underscore the need for additional features to distinguish signals of different biophysical origins. In particular, the amplitude of the pre-hyperpolarization peak and waveform polarity emerge as highly informative markers.

### Comparison across conditions

We first compared the two resting-state conditions. During eyes-closed periods, spiking was more tightly coupled to the ripple-band envelope elevations than during eyes-open rest (Supplementary Figures 3B, 5C), consistent with the emergence of globally synchronous up-states in this regime (Supplementary Figure 2C). The phase-locking preferences of the unit classes, however, were conserved across the two states (Supplementary Figures 3C, D). We hypothesize that the stronger coupling of spiking to the ripple band in the eyes-closed state reflects a network-wide recruitment of the oscillation-generating circuit, whereas the preferred phase, being set by the underlying wiring, remains conserved across states.

Turning to visual stimulation, when comparing unit activity during the elevated ripple band during the eyes-open rest with the activity evoked during the natural-image presentation (Supplementary Figure 5E, F), some specifics emerged as well. For the Biphasic and Wide units, NATIM was associated with increased activation and phase-locking. However, the preferred phases and relative timing between unit classes remained stable across conditions. This shift in activation points to a stronger activation of the sine-rhythm generating population in the evoked condition. In addition, the relatively strong up-modulation of the putative inhibitory NarrBI units in NATIM condition might reflect additional inhibitory influence specific for this regime. The Positive units, although being slightly more upmodulated in the NATIM condition, did not show increased phase locking (Supplementary Figure 5F), nor the sharp spiking transients (Supplementary Figure 8C). We hypothesize that the observed difference between the resting state and visually evoked unit activity possibly reflects distinct routing of information with inhibition acting as the router.

Overall, we conclude that although the strength of engagement differs between the conditions, high-frequency oscillatory activity remains tightly coupled to precisely organized spiking in all the described experiments.

### Limitations of the study

Our study has several limitations. First, the number of subjects is small in absolute terms: three macaque monkeys and two blind human volunteers. Three animals exceed the sample size of most non-human primate electrophysiology studies, and the reproduction of the main effects in human V1 provides independent cross-species confirmation, but between-subject variability was present.

Second, the temporal sampling of the resting state is sparse: seven sessions across three macaques, acquired at irregular intervals spanning up to a year of chronic implantation, with each recording day sorted independently and no tracking of units across days. We therefore cannot follow the recorded population continuously over the lifetime of the implant, nor separate genuine changes in circuit dynamics from slow displacement of the arrays and progressive lesioning of the surrounding tissue.

Third, we have no histological verification of electrode tip locations, as chronic lesioning of the tissue precluded it. The layer assignment rests on converging indirect evidence — shank length relative to cortical thickness, waveform priors from laminar Neuropixels recordings, spatial co-detection structure, and the alpha/gamma gradient — which agree with one another but are individually inconclusive.

### Future directions and clinical applicability

The insights derived from our data were made possible by state-of-the-art spike-sorting approaches^24^ and recent advances in waveform-based unit classification^19–21,36^, which enabled the reliable identification even of units with non-standard spike morphologies, such as opposite-polarity and triphasic. Our analysis motivates future work on the contributions of distinct cell types to oscillatory dynamics, particularly in non-human primates and in the human recordings. The ability to retrospectively infer laminar depth in Utah-array datasets and to isolate axonal activity may ultimately hold clinical relevance for stimulation strategies and brain–computer interface development, improving stimulation strategies by targeting specific layers or neuron types.

## STAR Methods

### Experimental model and study participant details

#### Macaque monkeys

We analyzed data from three macaque monkeys, as previously published in two datasets: monkey L from Chen et al. 2022 ^27^ and monkeys N and F from Papale et al. 2025, supplemented with additional resting-state recordings from N and F. Monkey A from Chen et al. 2022 was excluded due to an insufficient number of detectable single units, reflecting poor recording quality. Full details of surgical procedures and experimental setups are provided in the original publications. Briefly, each monkey was implanted with sixteen Utah arrays targeting visual areas V1, V4 (all monkeys), and IT (only in monkeys N and F), with exact placements varying across subjects. In monkey N, one array was removed intraoperatively due to technical issues. In the present study, we analyzed only V1 recordings: 14 arrays in monkey L, 7 in monkey N, and 8 in monkey F. Each Utah array comprised 64 electrodes (1.5 mm length) arranged in an 8 × 8 grid with 400 µm spacing. We utilize data from both the resting-state (RS) and natural-stimuli (NATIM) conditions. During RS sessions, monkeys were head-fixed in a room with no visual input and were allowed to remain awake or fall asleep. Monkey L contributed three RS recordings; monkeys N and F each contributed two. For monkey L, an eye-tracking camera was available, enabling validation of our eyes-open/eyes-closed classification procedure, which we subsequently applied to all subjects. For the NATIM condition, we analyzed the full recording sessions from Papale et al. 2025, processing the data agnostically with respect to trial structure and using the entire duration of each session.

#### Blind human volunteers

The dataset comprised two adult male volunteers with acquired blindness, with one 96-channel (1.5 mm length) Utah array implanted in V1 in each of the subjects. The Utah array was implanted in the subject’s right occipital cortex, near the occipital pole. The details of surgical procedures are described in Rocca et al., volunteers 2 and 3^40^. We considered the recordings of the resting state in the morning before the additional tasks and stimulations carried out during the day, and the last recordings of the resting state within an experimental day. The volunteers had the implant inserted for 6 months, after which it was explanted. In the first subject, the number of recordings was 94, with a total duration of 315.1 minutes. In the second subject, the number of recordings was 90, with a total duration of 224.9 minutes. Out of all the recordings, we are making 4 publicly available.

#### Registration and informed consent

Clinical trial registration at ClinicalTrials.gov, under identifier NCT02983370. The clinical trial protocol can be accessed at: https://clinicaltrials.gov/ct2/show/NCT02983370. The human experimentation was performed under a protocol that was approved by the Hospital General Universitario de Elche Clinical Research Committee and registered at ClinicalTrials.gov (NCT02983370). We followed all relevant ethical guidelines related to clinical trials regulation (EU No. 536/2014 (repealing Directive 2001/20/EC), the Declaration of Helsinki and the EU Commission Directives 2005/28/EC and 2003/94/EC), and we obtained written informed consent before any study procedure was conducted. All the procedures and risks were fully explained to the subjects prior to their participation, emphasizing the investigational nature of the study and that they should not expect any short- or long-term benefit resulting from participation in the study. A complete systemic, physical, neurological, and psychological evaluation was performed to assess the adequacy of the volunteers. They understood that the main purpose of the study was to gain knowledge essential for the future development of a cortical visual neuroprosthesis for the blind.

### Methods details

#### Preprocessing

Data from each channel and each recording day were processed and normalized independently, yielding single-unit spike trains, local field potentials (LFPs), ripple band (RB) signals, and delta-band signals. Because the source datasets differed in their original preprocessing pipelines, we did not rely on the preprocessing provided in the respective publications. Instead, we reprocessed all raw recordings, sampled at 30 kHz, using a unified pipeline applied consistently across all three monkeys and two human subjects. Most of the analysis code was implemented in Python.

##### Local field potential and delta band signal

To obtain the local field potential (LFP), the raw signal was first low-pass filtered at 250 Hz using a 6th-order Butterworth filter (*elephant.signal_processing.butter*) and then downsampled to 1 kHz with *scipy.signal.decimate*. The delta-band signal was obtained by applying an additional 0–4 Hz band-pass filter of the same type to the LFP. To extract the phase of the delta-filtered signal, we computed its analytic representation using *scipy.signal.hilbert* and derived the instantaneous phase from the angle of the analytic signal.

##### Ripple band

To extract the ripple band (RB) signal in the 80–150 Hz range, we band-pass filtered the raw 30 kHz data using a 6th-order Butterworth filter (*elephant.signal_processing.butter*) applied with *sosfiltfilt*. The filtered signal was then downsampled to 1 kHz using *scipy.signal.decimate*. We computed the analytic signal using *scipy.signal.hilbert*, from which we derived the RB envelope (via *np.abs*) and RB phase (via *np.angle*). Finally, we normalized the RB envelope by dividing it by its standard deviation for each channel.

##### Spike sorting

The raw data from all sessions were spike-sorted using a semi-automatic custom workflow combining *SpikeInterface*^47^ and *KiloSort4*^24^. First, we band-pass filtered the 30 kHz raw signals between 0.5 kHz and 6 kHz using a fourth-order Butterworth filter. The signals were then whitened to remove spurious correlations arising from electrode crosstalk, as described in Oberste-Frielinghaus et al. 2026^42^. The preprocessed signals were then passed to the Python implementation of *KiloSort4* for spike sorting. Briefly, *KiloSort4* detects spike events via template matching, clusters the resulting templates, and merges clusters based on auto- and cross-correlogram structure. We disabled all internal preprocessing steps in *KiloSort4*, particularly drift correction, because these procedures are optimized for acute laminar probes and produced undesirable artefacts when applied to two-dimensional Utah arrays.

After automatic spike sorting, each waveform cluster was evaluated against a fixed set of quality metrics. A unit was retained only if it satisfied all criteria simultaneously. Firing rate had to exceed 1 Hz, and this requirement was applied first because it also removes low-spike-count units for which the refractory and coincidence measures below cannot be computed. Waveform signal-to-noise ratio (SNR) was assessed using two complementary measures: the SpikeInterface SNR annotation (peak amplitude relative to the background noise of the continuous trace, threshold ≥ 2.0) and a metric derived directly from the sorted waveforms (template peak-to-peak amplitude divided by the standard deviation of each spike’s residual from the mean template, an isolation/consistency measure, threshold ≥ 4.0). Refractory-period contamination was quantified using a sliding violation ratio evaluated at a 1.5 ms window and required to remain below 1.5. Cross-unit coincidence was computed for every pair of simultaneously recorded units on an array as the fraction of spikes occurring within 0.5 ms of each other; each unit was assigned the ratio corresponding to its most coincident partner, which had to be below 10. Presence ratio (the fraction of binned recording-duration bins containing at least one spike) was required to exceed 0.75. Finally, locking to 50 Hz and 60 Hz electrical grid noise had to remain below 0.7. We processed the data from each day separately and did not track the same units across days. The results were stored in *NIX* format together with relevant metadata, including electrode positions, cortical area, and array identity. For the details on implementation and quality metrics see preprocessing code enclosed alongside the dataset.

##### Eyes open and closed classification

Eye closure in monkey L was monitored using an infrared camera with automated pupil detection (*Thomas Recording GmbH*). Details of the eye-tracking signal and processing steps are provided in Chen et al. 2022. For monkeys N and F, no camera-based eye tracking was available. In these cases, we inferred eye-open and eye-closed periods from changes in low-frequency LFP power (<12 Hz). We computed a full spectrogram of the LFP using a 1-s moving window and normalized the power within each frequency. We summed the power of all frequencies below 12 Hz, smoothed this signal with a 10-point moving average, and set an empirically determined threshold to split the eyes-open and eyes-closed periods. This procedure was validated against the camera-based measurements in monkey L, where ground-truth eye-tracking data were available^43^.

##### Noise channels identification

Channels exhibiting prominent line-noise contamination within the ripple band were excluded using the following procedure. From the ripple-band–filtered signal, we computed the average spectral power in the 90–95 Hz and 98–102 Hz intervals using *scipy.signal.welch*. If the mean power in the higher interval (98–102 Hz) exceeded that of the lower interval, the channel (and all single units recorded on it) was excluded from further analysis. An analogous procedure was applied to identify contamination at 120 Hz, using the 110–115 Hz and 118–122 Hz intervals.

#### Ripple detection

Ripple events were detected using the dual-threshold algorithm (*neurodsp.burst.detect_bursts_dual_threshold*; as described in Feingold et al. 2015). Briefly, this method uses two thresholds [A, B], A<B, and identifies intervals during which the maximum of the signal envelope exceeds B while the minimum remains above A. In our case, the bursts were detected on the RB envelope smoothed with 40-Hz lowpass filter, using thresholds of [2.5, 3.5] of standard deviations of the envelope. Burst duration was defined as the interval between crossings of the lower threshold A. Each recording day and each channel were processed independently. Bursts shorter than 40 ms were discarded. In the macaque analysis, after detection on the full recording, ripples were assigned as eyes-closed (EC) if they fully fell into the EC interval, or eyes-open otherwise. For the Natural Images dataset, no further subdivision into trials or stimulus periods was applied; the entire session was analysed continuously.

#### Ripple properties calculation

The ripple rate for each channel was defined as the number of detected ripples per second, computed separately for each recording session. For resting state data, we further refined this metric by calculating eyes-open and eyes-closed ripple rates using only the ripples and time intervals corresponding to the respective eye-state classification described above. Ripple frequency was estimated as the number of ripple band cycles between the first and last zero-crossings of the RB signal within the event, divided by the duration between those crossings. Ripple amplitude was defined as the maximum value of the ripple band envelope during the event.

#### SUA classification

For every single unit, we computed an average waveform by taking the mean of all spike waveforms returned by the spike-sorting algorithm. This average waveform was then z-scored, and we refer to the resulting normalized trace as the unit’s NormAWF. Units were classified as negatively dominant if the absolute value of the global minimum of the NormAWF exceeded the global maximum. Otherwise, they were labelled positively dominant. For each unit, the height of the first peak was defined as the global maximum occurring before the global minimum of the NormAWF. Waveform width was defined as the time difference between the global minimum and the subsequent global maximum, converted to microseconds. Among negatively dominant units, we distinguished three classes based on waveform width: narrow (≤ 235 µs), medium (235–430 µs), and wide (> 430 µs). Within the narrow and medium groups, we further separated units into biphasic and triphasic subclasses, with biphasic units defined as having a first-peak height below 1.2 z-scored amplitude units, and triphasic units comprising all remaining cases.

#### SUA properties calculation

Because of the recording setup geometry, each unit was assigned only to a single electrode. The firing rate of each unit was defined as the number of spikes per second over the full duration of the recording. To assess how firing rates were modulated by the RB envelope, we computed the ratio between the firing rate during periods when the RB envelope on that channel exceeded its median and the firing rate during periods when it fell below the median.

##### Phase selectivity

For each unit, we considered the ripple band phase sampled at 1 kHz from the electrode on which the unit was recorded (see Preprocessing). Spikes were binned into 1-ms bins, and we extracted the RB phase values corresponding to bins containing at least one spike. To compute the normalized RB phase selectivity and the preferred RB phase of each unit, the spike-associated phases were binned into a 25-bin histogram spanning the interval (− π, π] and normalized to probability density. We then computed the circular mean of the set of vectors defined by the bin-center angles and their corresponding densities. The angle of the resulting vector was taken as the preferred phase ∈ (− π, π], and its length served as the normalized phase-selectivity index ∈ [0, 1].

##### Correlation of spikes and the elevated ripple band in RS

For each recording and a given array, we summed the spiking activity across all units on the array, and also computed the sum of the RB envelope for the array. The data were segmented into 1-s windows, each labelled as eyes-open (EO) or eyes-closed (EC) based on the classification procedure described above. We then computed the Pearson correlation between the array-level spike count and RB envelope sum using 10-ms binned versions of the labelled data vectors.

##### Graph of the spatial clustering of single-unit types in RS

We quantified spatial clustering between pairs of single-unit classes as follows. For each class combination, we calculated N_close, the number of unit pairs recorded either on the same channel or on one of the eight adjacent channels (differing by at most one row and one column on the array grid) within the same array, considering only units located in areas V1. Statistics were aggregated across all resting-state (RS) and naturalistic-image-viewing (NATIM) sessions, and across all recording days and animals.

To evaluate whether the observed co-occurrence exceeded chance, N_close was compared to a null distribution obtained by global permutation: within each recording type, unit-class labels were randomly shuffled 1000 times across all units pooled over arrays, dates, and animals, preserving the overall class composition and channel adjacency while allowing class identity to be freely exchanged between arrays and animals. The RS and NATIM null distributions were then combined permutation-by-permutation. The expected co-occurrence for each class pair, N_exp, was defined as the mean of the 1000 pooled permuted counts.

The clustering (enrichment) ratio was computed as N_close/N_exp: a value of 1 indicates chance-level co-occurrence, values above 1 reflect spatial enrichment, and values below 1 indicate spatial avoidance. Enrichment ratios are visualized as color-coded edges in Figure 5A (blue: depletion; red: enrichment; 1 = chance), with the color scale fixed to 0.25–1.5 and values outside this range clipped to the endpoint color to prevent any extreme pair from compressing the scale. No significance threshold was applied; only the enrichment ratio is represented.

##### Assignment of channels to the two clique-defined unit groups

For each channel, we assessed whether it contained more single units belonging to the blue-clique unit group (putative Deep layer, containing Triphasic + Positive units) or the orange-clique unit group (putative Layer 4, containing Biphasic units) cell types, as defined in Figure 5A. Channels with no units, or with equal proportions of units from both clique groups, were left unclassified. We pooled units across all recording days, all conditions, per animal.

#### Hypnogram

Hypnogram estimation of the macaque resting state data for a given array and recording day was done using *yasa.SleepStaging* automatic sleep classification algorithm ^44^. For each channel, the LFP signal served as the sole input, no EOG nor EMG channels were provided. The algorithm returns, for every 30-s epoch, the probability that the segment corresponds to wakefulness, REM sleep, or one of the three NREM stages (N1, N2, N3). To obtain an area-level confidence estimate for a given sleep stage S during the time window T, we average the probabilities of each channel being in stage S in the time window T, across all the channels in the cortical area.

#### Triggered statistics

##### Ripple-triggered LFP

For each channel, we used the positive peaks of the detected ripples (restricted to EO or EC periods in macaque resting-state data) as trigger events. Around each trigger, we extracted and averaged the z-scored LFP within a predefined time window. For the final plots (Figure 5E, Supplementary Figure 8C, 9F), these per-channel averages were further averaged across all recordings and all animals/volunteers.

##### Ripple-triggered wavelet spectrograms in RS

We computed a wavelet spectrogram from the LFP within a 1-s window centred on the ripple’s maximal positive peak, using complex Morlet wavelets (*cmor1.5–1.0, PyWavelets*) with 180 logarithmically spaced scales between 0.4 and 3. For each channel and recording, we averaged the resulting time–frequency representations across all ripples of a given type (EC or EO). For visualisation, these per-recording spectrograms were further averaged across channels and animals, and each frequency row of the final spectrogram was z-scored to account for the 1/f spectral background.

##### Aligned ripple-triggered phase

For each ripple, we identified the first trough of the RB-filtered signal following the ripple onset. This trough was labeled as phase 0, and the RB phase within a window of N seconds around this point was realigned accordingly. Then, for each unit recorded on the same channel, we collected all spikes occurring within N seconds of the first trough and stored the corresponding realigned RB phase at spike time. Data from all units and all ripples were then pooled, for either all resting-state (RS) or natural-stimuli (NATIM) recordings, to construct a histogram of spike-aligned phases for each spike-class category. For visualisation, the resulting histograms were smoothed using a 1-ms Gaussian kernel.

##### Delta-trough-triggered delta and ripple band envelope in RS

All analyses were performed at the array level. First, we computed the sum of the ripple band envelopes across all channels on the array, as well as the sum of the delta-band (0–4 Hz) filtered signals. Troughs of the summed delta signal (detected by *scipy.signal.find_peaks()* with default settings) served as trigger events. Using these triggers, we computed the average ripple band envelope and the average delta-band signal using the previously summed array-level activity.

#### Computational model of Positive units population activity

We modelled the activity of the positively dominant unit population using the NEST simulator ^30^. The network consisted of 2500 current-based leaky integrate-and-fire neurons (*iaf_psc_alpha*) without mutual connections. The neurons were instantiated with the following parameters.

| Parameter | Value | Parameter | Value |
| --- | --- | --- | --- |
| C_m | 250 pF | E_L | -70 mV |
| tau_m | 10 ms | V_th | -55 mV |
| t_ref | 2 ms | V_reset | -70 mV |
| tau_syn_ex | 2 ms |  |  |

As the input for the population, we used 3200 sinusoidally modulated Poisson spike generators (*sinusoidal_poisson_generator*). Each generator produced spikes with a baseline rate of 80 Hz, and sinusoidal modulation of amplitude 30. The modulating sine had a frequency of 100 Hz, and the initial phases of the sine wave were picked from the Gaussian distribution with 0 mean and a standard deviation of 0.05*2π. The spike generators were connected to the neuronal population with the *pairwise_bernoulli* rule with 0.05 connection probability, weight 5, and delay 1.

### Quantification and statistical analysis

#### Phase selectivity

To estimate a shuffle distribution for normalized selectivity and determine a significance threshold, we used the 1-ms binned spike train together with the RB phase vector for the full recording session. For each unit, the spike train was kept fixed, while 100 circularly shifted versions of the phase vector were generated using *np.roll*, with shift values drawn uniformly from *[0, recording duration]*. For each shifted phase vector, normalized phase selectivity was computed as described above. Shuffle-derived selectivity values were pooled across all units detected in the macaque RS recordings from all animals and binned into a 100-bin histogram over [0,1]. The threshold for classifying a unit as phase-selective was defined as the 99th percentile of this shuffle distribution rounded up to two decimal places (0.04). The same value of threshold was used in the NATIM and human datasets, for comparability.

#### Statistical testing

Statistical tests were presented in Figure 1K (testing whether the correlation is negative), Figure 5C,D (comparing Alpha and Gamma in clustered layout in macaque EO RS), and in Supplementary materials: Supplementary Figure 3B (testing spikes and RB envelope correlation between RS EC and RS EO), Supplementary Figure 3C, D (comparison of EC and EO properties per unit class), Supplementary Figure 5E,F (comparing single-unit properties in RS EO and NATIM), Supplementary Figure 8A, B (comparing Alpha and Gamma in clustered layout in macaque NATIM).

The exact type of the test and p-value are reported in Figure captions. All statistical analyses were performed using the Python *scipy.stats* module, and can be found in the Jupyter notebooks for the respective figures. The number of data points used in each of the analyses was the following.

Figure 1K (one-sided Wilcoxon test): n=72 windows where the Pearson correlation was computed.

Figure 5C,D (one-sided Mann-Whitney U-test): n_blue_L=202, n_orange_L=328, n_blue_F=116, n_orange_F=140, one data point represents the spectral power in the given band, on one channel in one recording day, per animal, RS macaque recordings, only EO periods considered. The number of data points is the same for Alpha and Gamma bands. Unclassified channels were not included in the analysis.

Supplementary Figure 3B (two-sided Mann-Whitney U-test): n_EC=43341, n_EO=37714, one data point is a windowed Pearson correlation coefficient representing one Utah array, 1-s time bin of 10-ms binned spikes, EC or EO period only in a given recording. The number of bins in EC and EO differ, thus the number of data points is different. Pooled across monkeys and across recording days.

Supplementary Figure 3C, D (two-sided Mann-Whitney U-test, Bonferroni corrected): One data point represents one single unit of a given class. Number of units per class in EC and EO (the same): n_NarrBI=214, n_MedBI=527, n_Wide=462, n_NarrTRI=157, n_MedTRI=179, n_Pos=69. The number of units in Figures 2G, 3B, C is also the same (visually depicted in Figure 2F).

Supplementary Figure 5E, F (two-sided Mann-Whitney U-test, Bonferroni corrected): One data point represents one single unit of a given class. Number of units per class in NATIM: n_NarrBI=1398, n_MedBI=2029, n_Wide=2799, n_NarrTRI=1431, n_MedTRI=1058, n_Pos=659. The number of units in NATIM in Figures 4C, D, is also the same (visually depicted in Figure 4B). Numbers of points in EO as in Supplementary Figure 3C, D, as listed above.

Supplementary Figure 8A, B (one-sided Mann-Whitney U-test): n_blue_N=2419, n_orange_N=2488, n_blue_F=1375, n_orange_F=1567. One data point represents the spectral power in the given band, on one channel in one recording day, per animal, NATIM macaque recordings. The number of data points is the same for Alpha and Gamma bands. Unclassified channels were not included in the analysis.

## Declaration of generative AI and AI-assisted technologies in the writing process

During the preparation of this work, the authors used Claude to assist with rephrasing and editing of text, and for modification of analysis code, and Consensus for literature search. After using these tools, the authors reviewed and edited the content as needed and they take full responsibility for the content of the published article.

## Acknowledgements

We would like to thank Sanja Bauer Mikulovic (Medical University of Vienna, Austria), Ján Antolík (Charles University, Czechia) and the Computational Systems Neuroscience Group at the Charles University in Prague for fruitful discussions and constructive feedback. The work was supported by the Charles University grant PRIMUS/24/MED/007; Charles University Grant Agency grant (GAUK) No. 224226; NWO (Dutch Research Council) grant VENI No. VI.Veni.222.217; ERC Starting grant “SteerMEM”, No. 101219949; Programme Johannes Amos Comenius (OP JAK) under the project ’MSCA Fellowships CZ - UK3’, reg. n. CZ.02.01.01/00/22_010/0008220; NWO (Dutch Research Council) Crossover grant No. 17619 “INTENSE”; Spanish Ministerio de Ciencia, Innovación y Universidades / Agencia Estatal de Investigación grants PDC2022-133952-I00 and PID2022-141606OB-I00; MICIU/AEI/10.13039/501100011033 grant AIA2025-164108-C31; Generalitat Valenciana grant CIPROM/2023/25; CYTED grant NT4SM; European Union’s Horizon 2020 Research and Innovation Programme under Grant Agreement No. 899287 (NeuraViPeR).

## Author’s contributions

Conceptualization, K.S. and K.K.; Methodology, K.S. and K.K.; Software, K.S. and A.M.-G.; Formal Analysis, K.S.; Data Curation, K.S. and A.M.-G.; Investigation, A.L., X.C., P.P., C.S.-S., and E.F.; Resources, A.L., X.C., P.P., C.S.-S., and E.F.; Visualization, K.S.; Writing – Original Draft, K.S. and K.K.; Writing – Review & Editing, all authors; Supervision, K.K., E.F. and P.P.; Funding Acquisition, all authors.

## Code and Data Availability

Code: NEAM-MFF/Ripples_project: Cortical ripples in the macaque V1 data analysis. Data: https://doi.org/10.12751/g-node.xg19b5

## Supplementary

**Supplementary Fig. 1.**
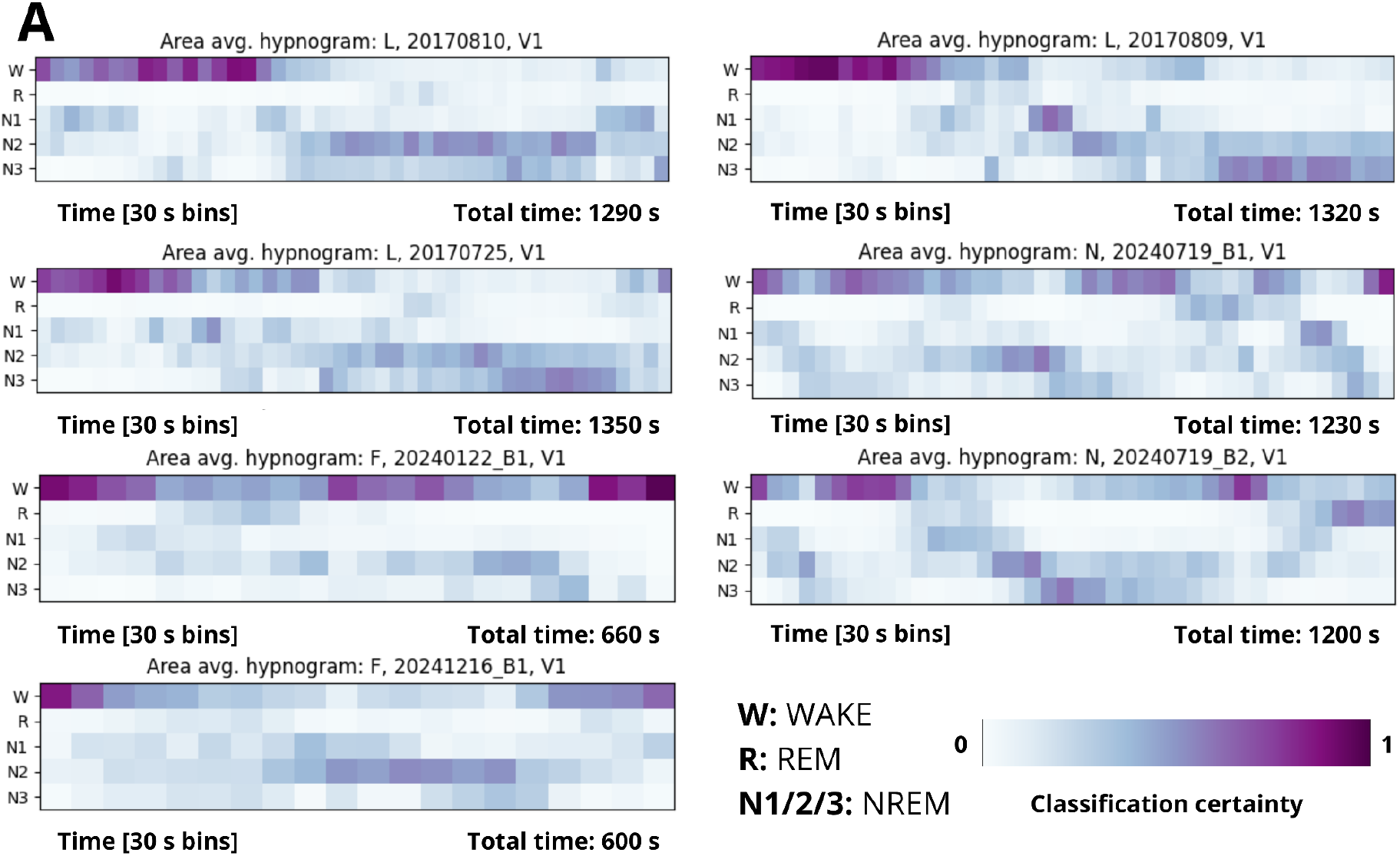
(Hypnograms and Wavelet spectrograms. Resting state.) **A.** Hypnograms for each recording session, calculated using all channels in V1. An automated sleep-scoring algorithm YASA was used.

**Supplementary Fig. 2.**
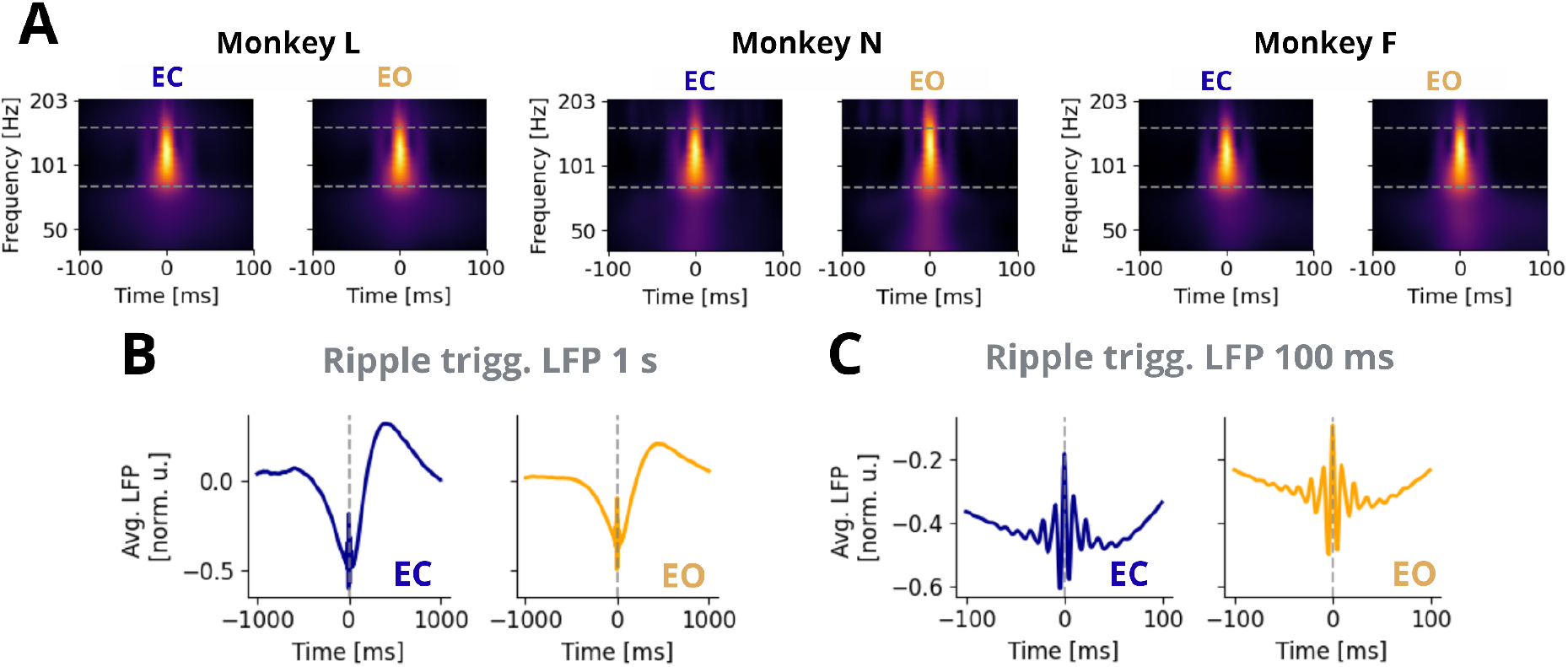
(Slow phenomena.) **A.** Ripple peak triggered wavelet spectrogram, separate for eyes closed (EC) and open (EO) conditions and per animal. Rows are z-scored to compensate for the 1/f spectral component. Figure 1D shows data pooled across animals. **B., C.** Average ripple-triggered LFP on a 1-s and 100-ms scale.

**Supplementary Fig. 3.**
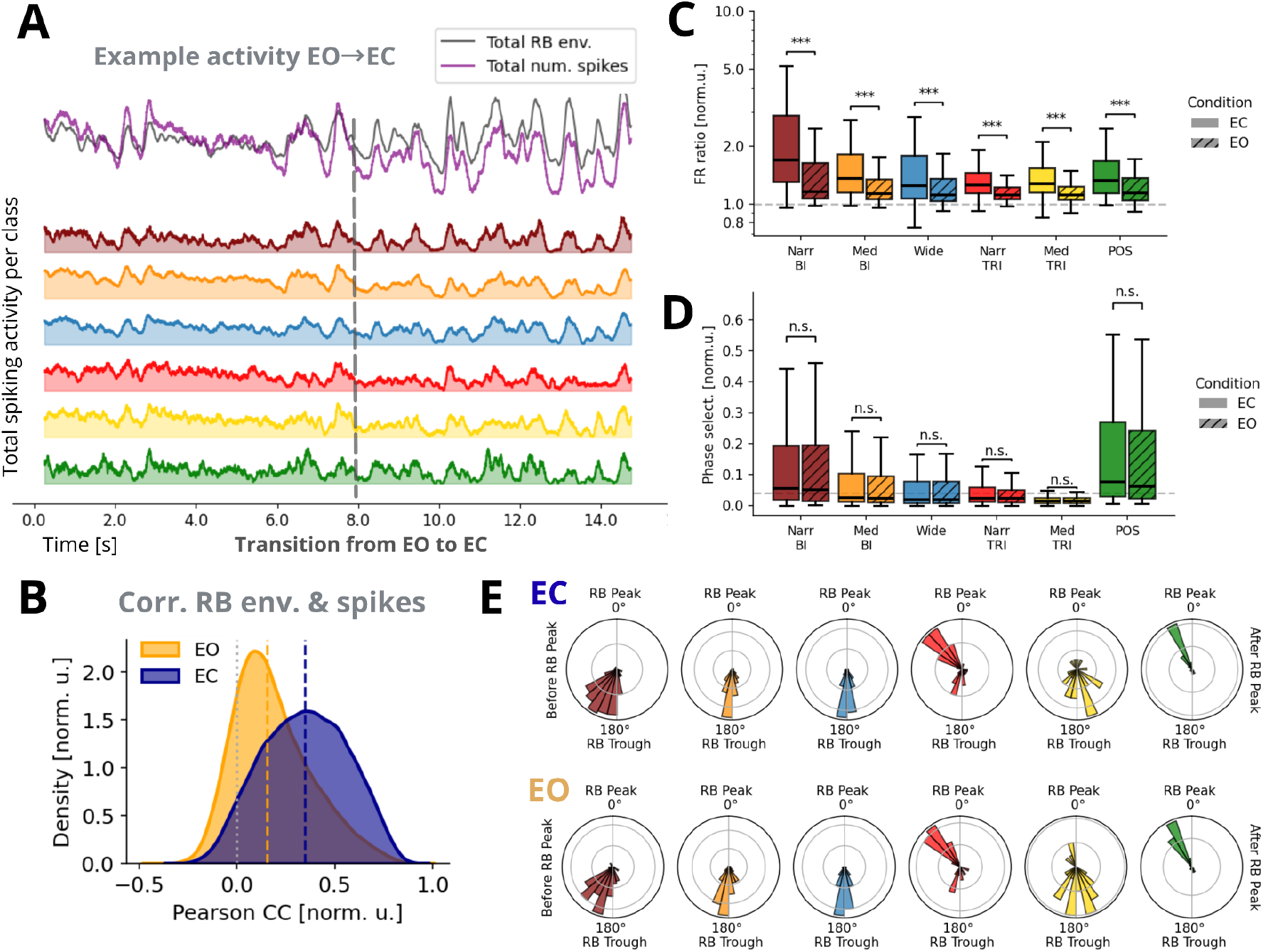
(Resting state, eyes closed/open comparison.) **A.** Example 15 s of activity, transition from EO to EC, monkey L, same time period as in Supplementary Figure 2C. Total spiking activity per unit class is color-coded. The sum of the spiking activity across the classes is shown as a purple line. The total RB envelope across all the channels is in dark gray. **B.** The distribution of the Pearson correlation coefficient between the total spiking activity and the total RB envelope per array. Dashed lines show medians. Data were binned into 10-ms windows, then split into 1-s moving windows, and a correlation coefficient was calculated in each window, further labeled as EC and EO. Data were pooled across all arrays. **C.** Ratios of firing rates during the RB envelope above its median value, and below its median value. Data during the resting state EC were tested against the resting state EO. All boxplots display the median and the interquartile range. Mann-Whitney two-sided U test, Bonferroni-corrected P-values: p<<10^-3^ for all classes. **D.** Normalized RB phase selectivity for a given unit class. Shuffle-based threshold in dashed gray. Data during the resting state EC were tested against the resting state EO. Mann-Whitney two-sided U test. Bonferroni-corrected P-values: p=1 for all unit classes. **E.** The histogram of preferred RB phases of selective units. One data point corresponds to one single unit of a given type. RS EC top plot, RS EO bottom.

**Supplementary Fig. 4.**
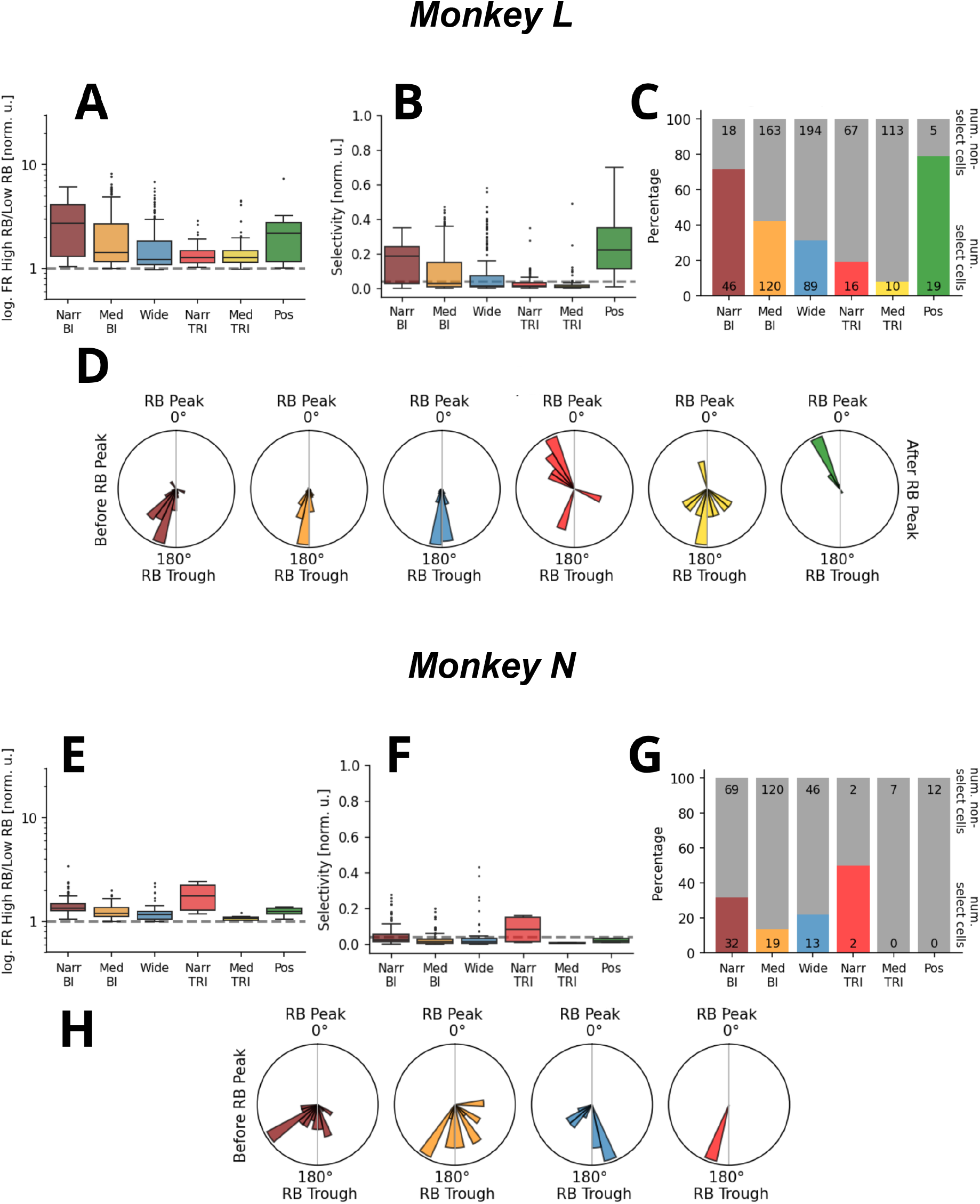
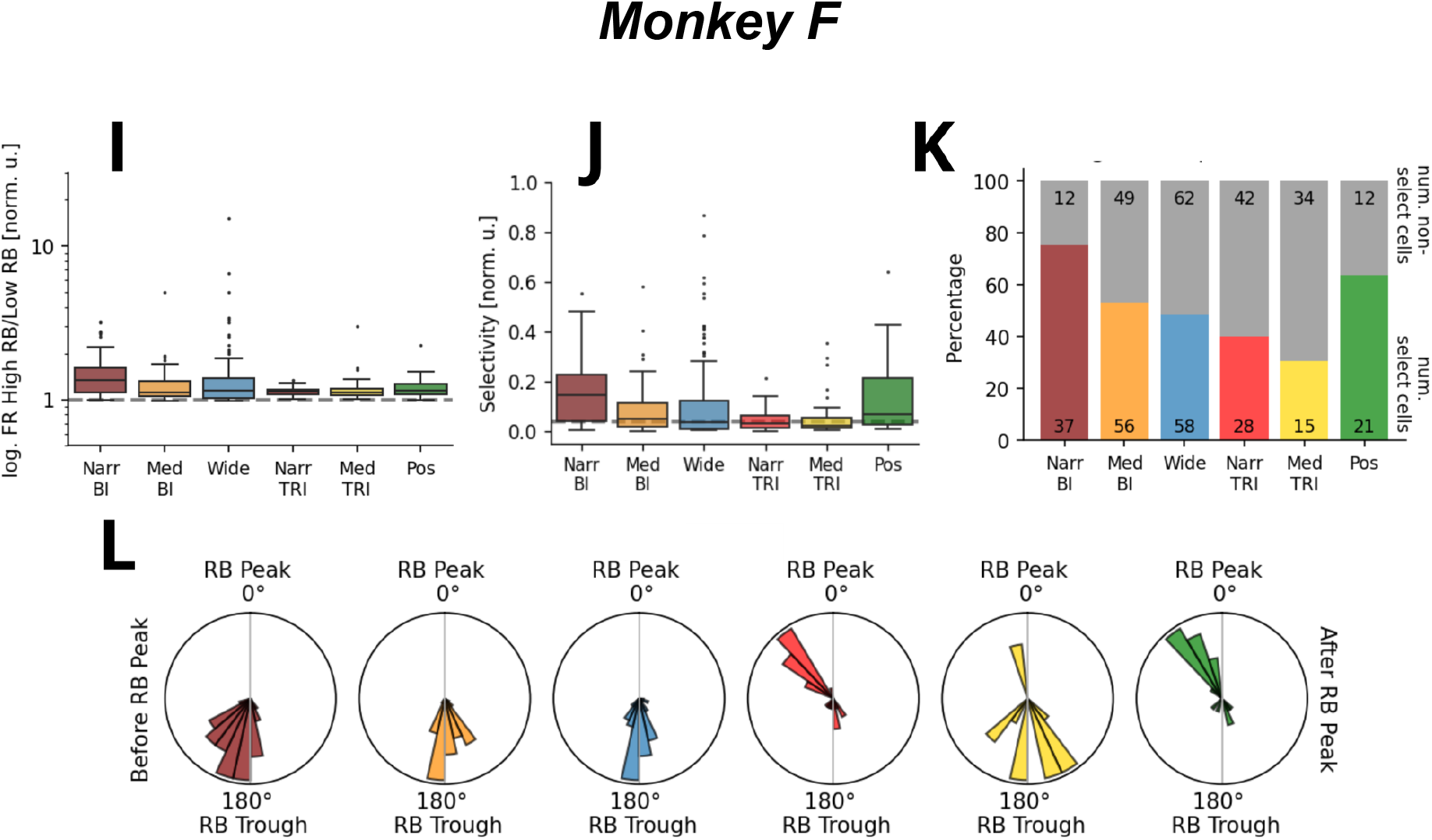
(L, F, N, Resting state, individual animals.) Overview of resting state statistics for individual animals. References point to the Figures in the Results. **A, E, I.** Ratio of firing rates during the RB envelope above its median value, and below its median value. Pooled per unit class, log-scale. As in 3B. **B, F, J.** Normalized RB phase selectivity for a unit class. Shuffle-based threshold in dashed gray. As in 3C. **C, G, K.** Number and ratio of RB phase selective units. As in 3D. **D, H, L.** The histogram of preferred RB phases of selective units. One data point corresponds to one single unit of a given type. As in 3E.

**Supplementary Fig. 5.**
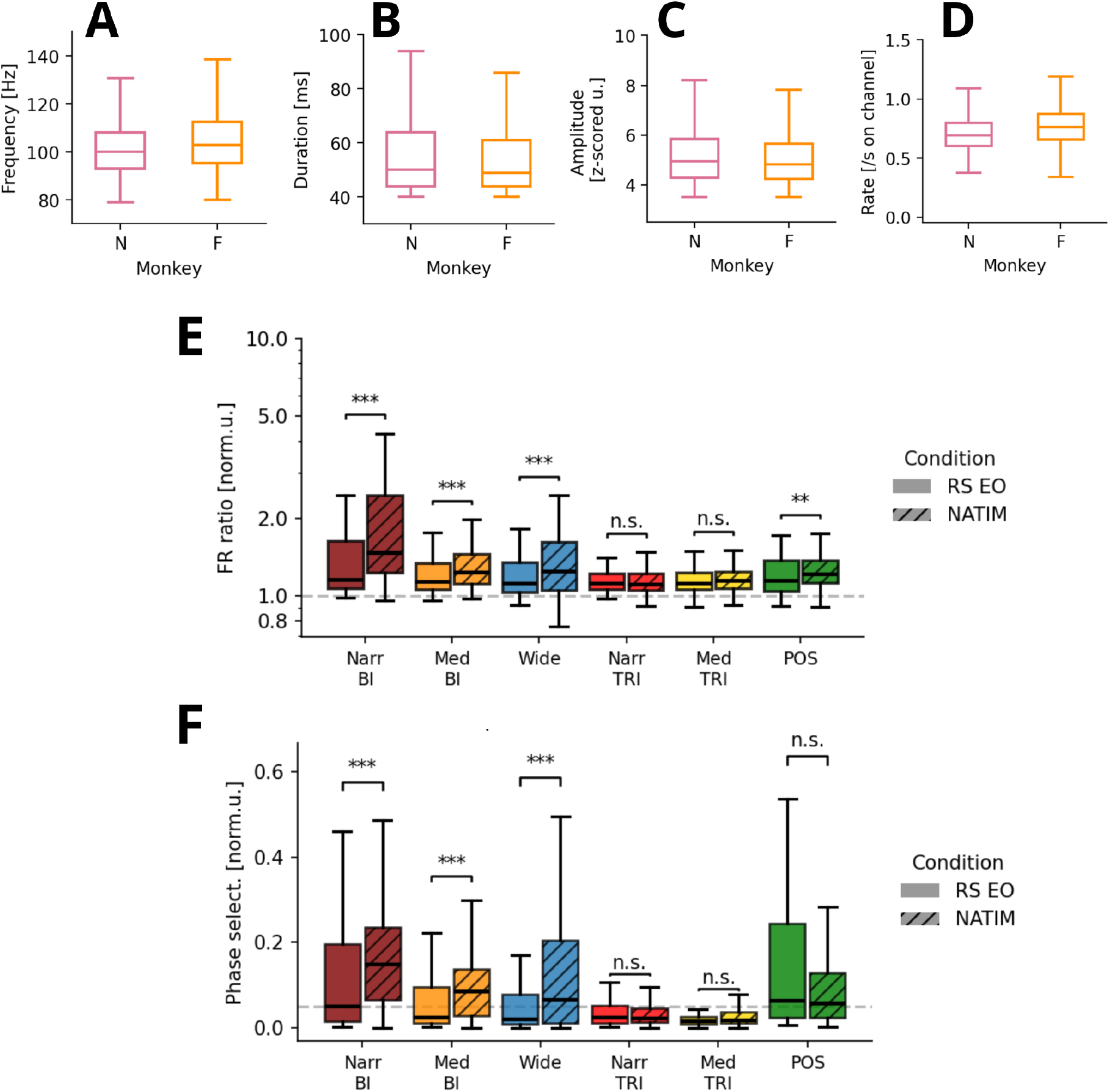
(Ripples in NATIM, comparison NATIM and RS EO.) Ripple detection in the Natural images dataset. Boxplots show median and interquartile range. **A, B, C.** Boxplots of the frequency of ripples, their duration, and amplitude. One datapoint corresponds to one ripple. Pooled via all channels, all recordings, per animal. **D.** Number of ripples per second. One datapoint corresponds to one channel during one recording day. **E.** Ratios of firing rates during the RB envelope above its median value, and below its median value. Data during the RS EO were tested against the natural images stimulation, i.e., Supplementary Figure 3C against Figure 4C. All boxplots display the median and the interquartile range. Mann-Whitney two-sided test. P-values per unit class, Bonferroni corrected: NarrBI, MedBI, Wide: p<<10^-5^, NarrTRI: p=1.0, MedTRI: p=0.36, Pos: p=8.6*10^-3^. **F.** Normalized RB phase selectivity for a unit class. Shuffle-based threshold in dashed gray. Data during the RS EO were tested against the natural images stimulation, i.e., Supplementary Figure 3D against Figure 4D. Mann-Whitney two-sided test, Bonferroni corrected. P-values per unit class: NarrBI, MedBI, Wide: p<<10^-5^, NarrTRI, Pos: p=1.0, MedTRI: p=0.20.

**Supplementary Fig. 6.**
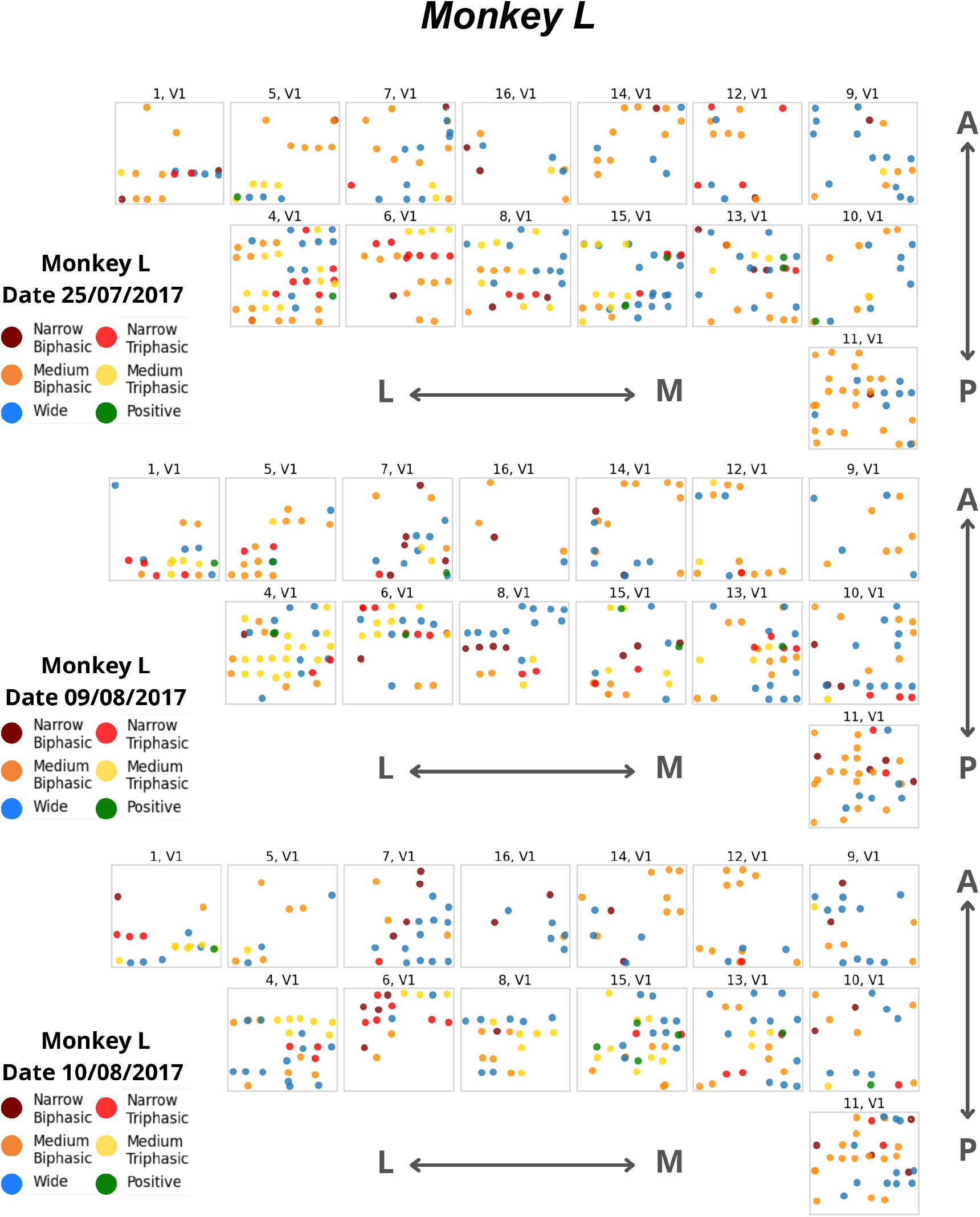
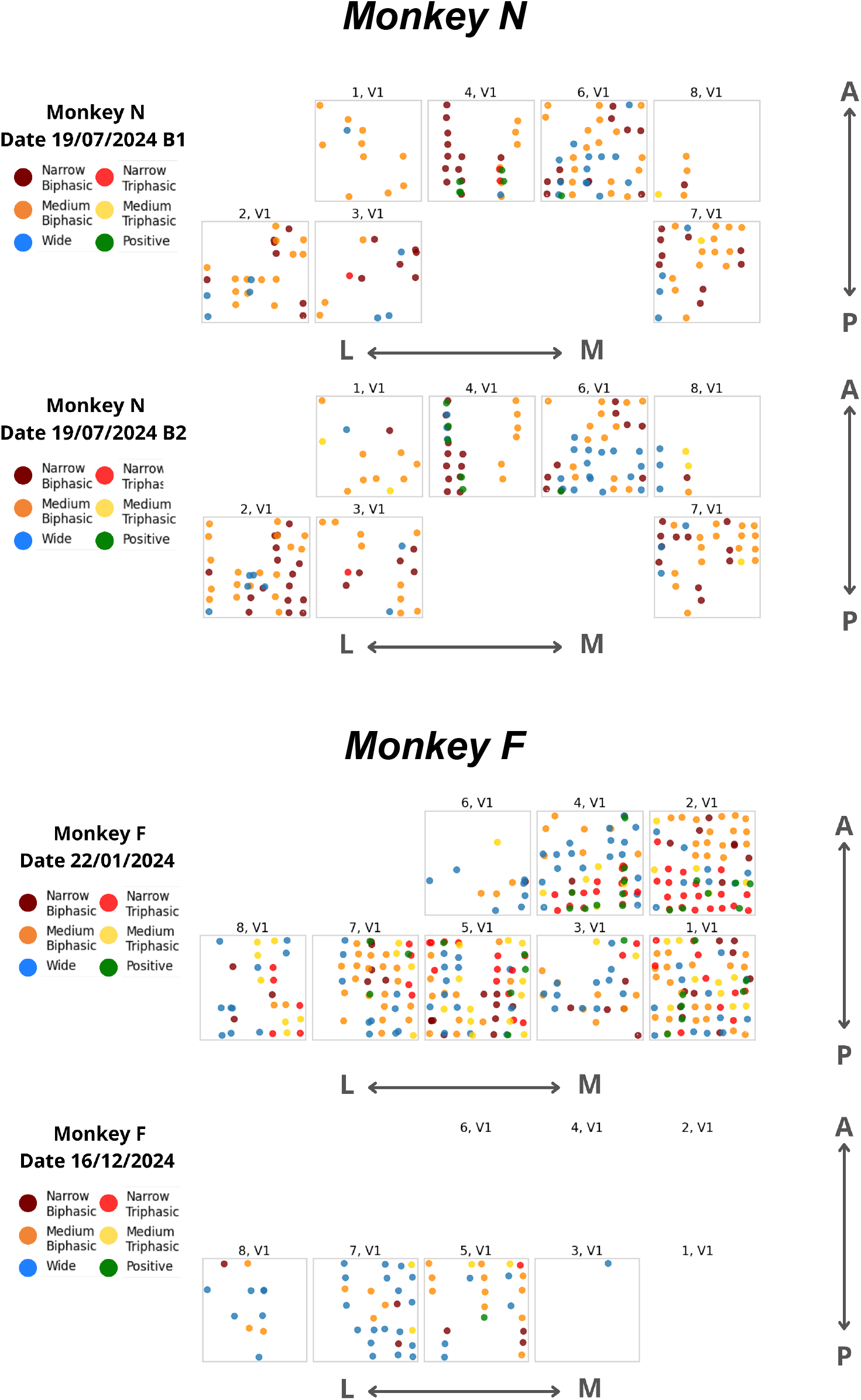
(Single units. Resting state.) Single unit distribution in the approximate cortical coordinates, one figure per recording. Gray boxes depict Utah arrays; titles indicate the array ID number. Types of units are color-coded in accordance with Figure 2. Slight jitter is added to the position of the channel on which the given unit was detected to accommodate multiple units co-detected on the same channel. Anterior-Posterior and Medial-Lateral orientations shown by arrows. In the second recording of monkey F, there were arrays with no units detected, thus not plotted.

**Supplementary Fig. 7.**
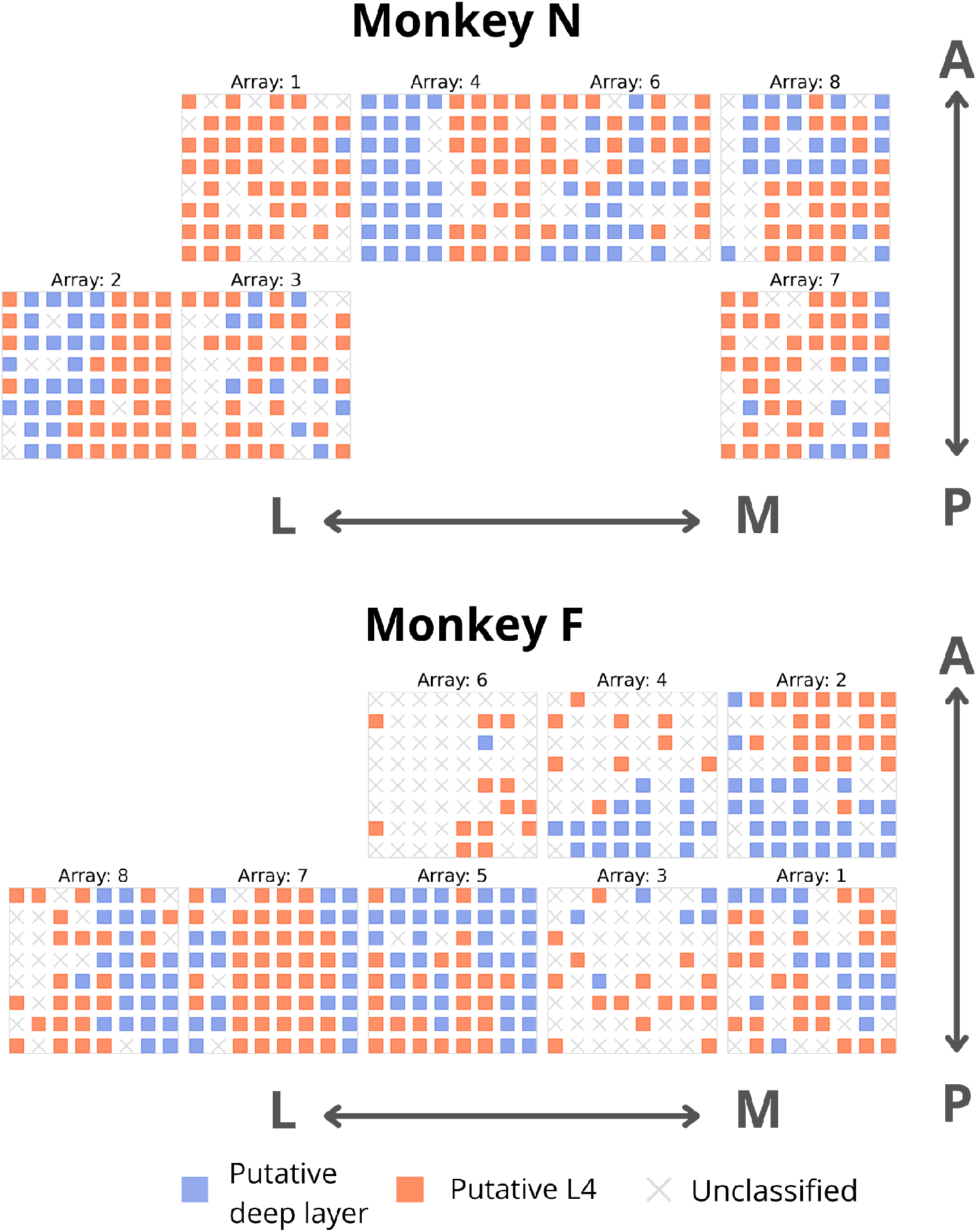
(Channel assignments to clusters. NATIM + RS units pooled.) For each channel, we assessed whether it contained more single units in the blue-cluster unit group in Figure 5A (NarrTRI+MedTRI+Pos) or the orange-edge unit group (NarrBI+MedBI). Channels with no units, or with equal proportions of units from both subgroups, were left unclassified. Pooled all RS and NATIM recordings per subject; monkey L is shown in main Figure 5B.

**Supplementary Fig. 8.**
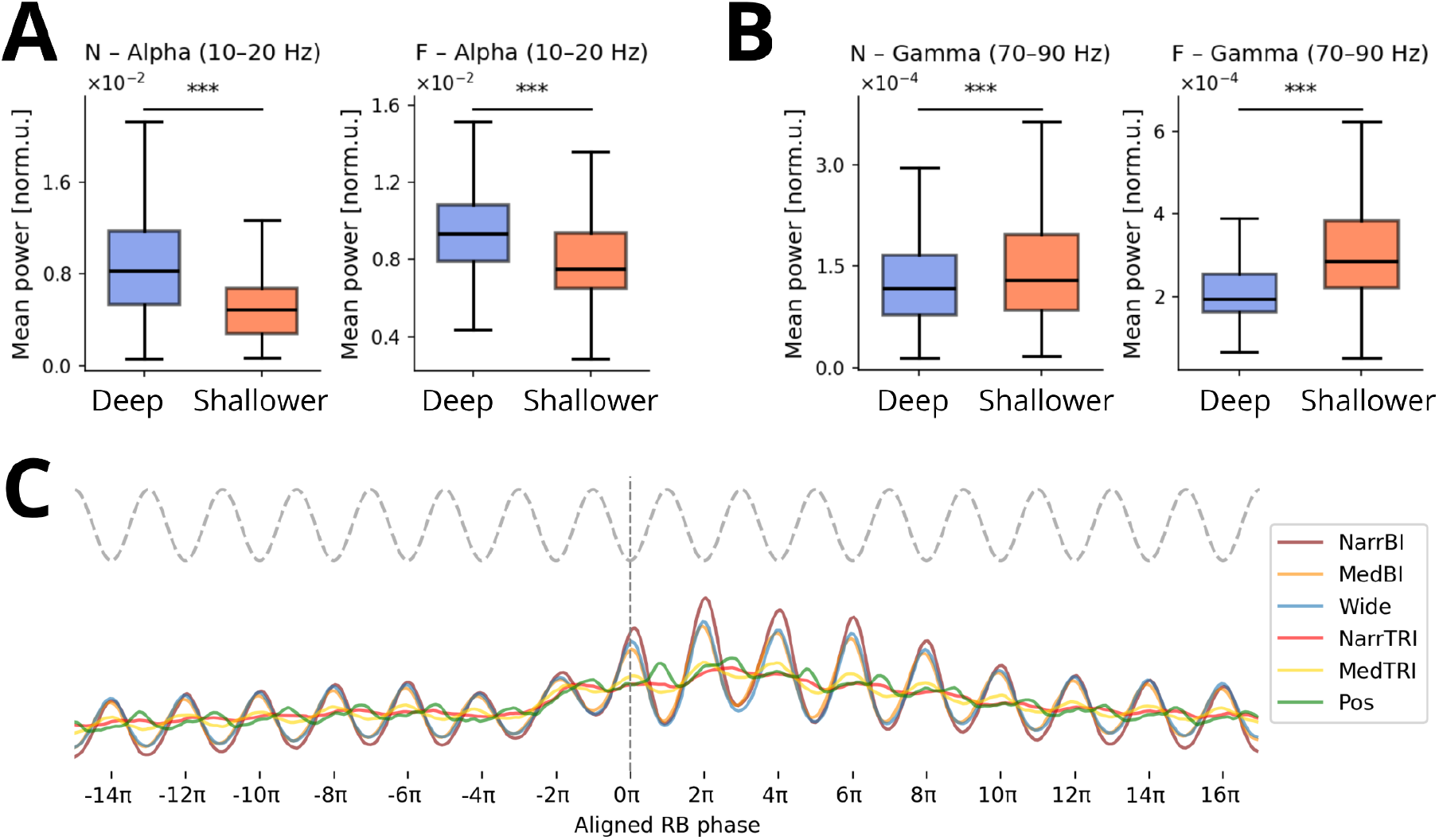
(Additional NATIM analyses.) **A.** Geometric means of power per channel in NATIM fixation periods in the Alpha 10-20Hz band, comparing putative deep-layer channels and putative shallower channels. **B.** Same as A., but for the Gamma 70-90Hz range. We tested with a one-sided Mann–Whitney test. For both monkeys, both frequency bands: p-val<<10^-5^. **C.** The first ripple oscillatory trough triggered phase-aligned spikes in NATIM. We aligned the spikes to the RB phase, and pooled the spiking activity around all the detected ripples. Phase 0 corresponds to the trigger. Both animals (monkeys N and F), all NATIM recordings pooled. Analogous plot for RS in Figure 5E.

**Supplementary Fig. 9.**
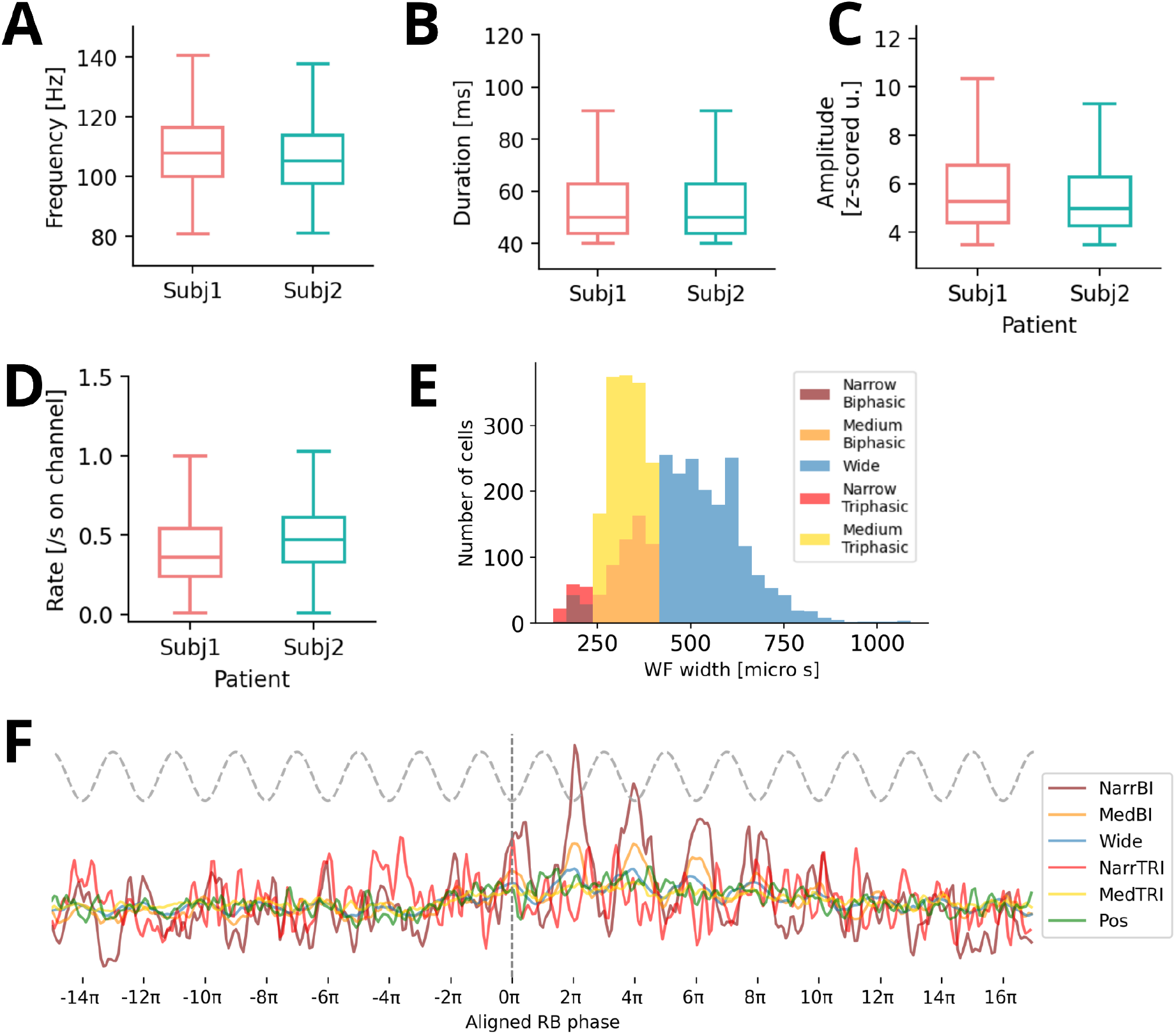
(Blind human comparative analysis.) Ripple detection and additional single-unit analysis in the blind human V1 dataset. Boxplots show median and interquartile range. **A, B, C.** The boxplots of the frequency of cortical ripples, their duration, and amplitude. One datapoint corresponds to one ripple. Pooled via all channels, all recordings, per subject. **D.** Number of ripples per second. One datapoint corresponds to one channel during one recording day. **E.** Distribution of the waveform widths W, only negative dominant waveforms included. Color-coded based on the classification criteria. As in Figure 2B. **F.** The first ripple oscillatory trough triggered phase-aligned spikes. We align the spikes to the RB phase, and pool the spiking activity around all the detected ripples. Phase 0 corresponds to the trigger. Pool across all subjects. As in Figure 5E.

**Supplementary Fig. 10.**
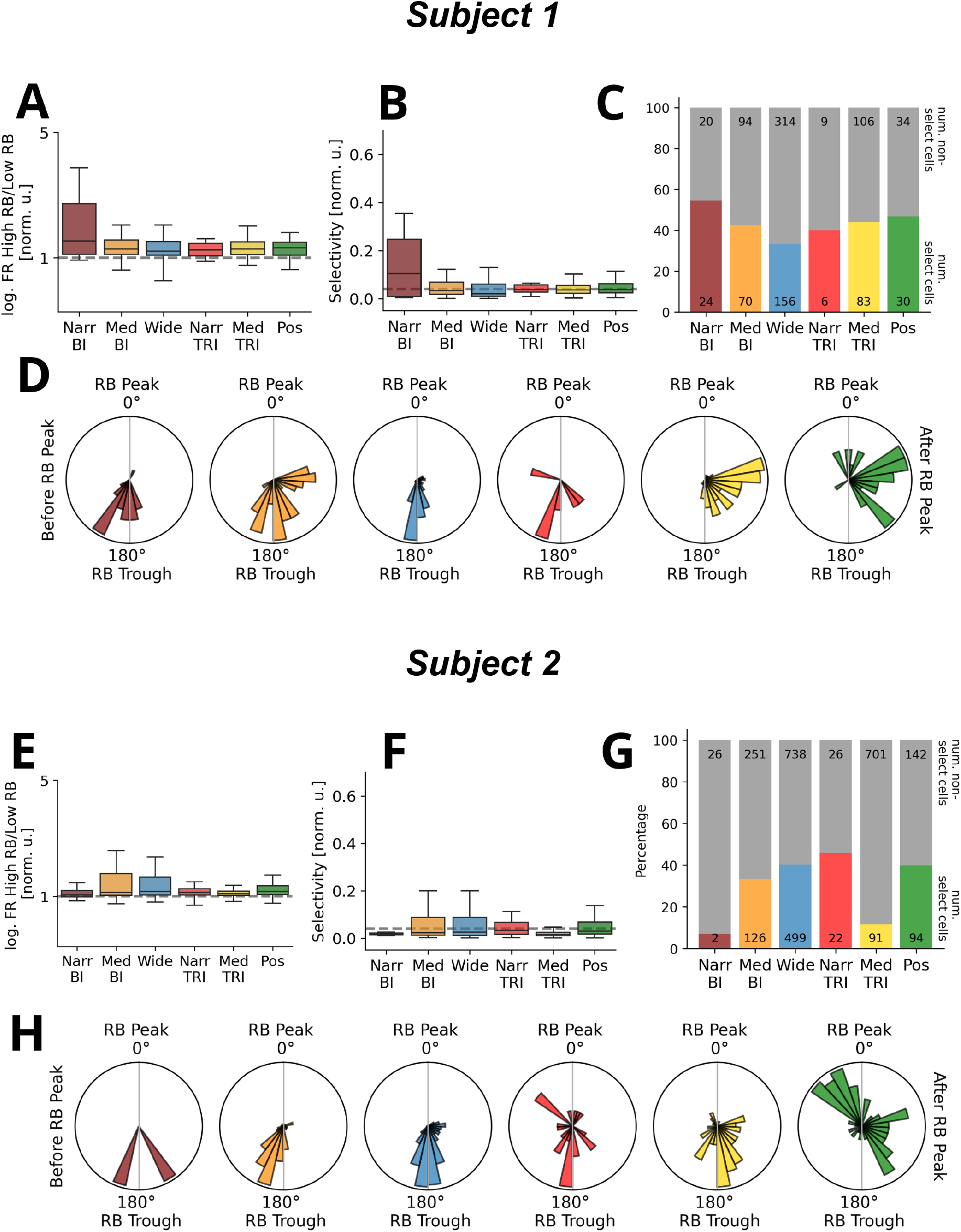
(Blind human, individual subjects.) Overview of resting state statistics for individual blind subjects. References point to the Figures in the macaque RS results. **A, E.** Ratio of firing rates during the RB envelope above its median value, and below its median value. Pooled per unit class, log-scale. As in 3B. **B, F.** Normalized RB phase selectivity for a unit class. Shuffle-based threshold in dashed gray. As in 3C. **C, G.** Number and ratio of RB phase-selective units. As in 3D. **D, H.** The histogram of preferred RB phases of selective units. One data point corresponds to a single unit of a given type. As in 3E.

